# Dynamic evolution of EZHIP, an inhibitor of the Polycomb Repressive Complex 2 in mammals

**DOI:** 10.64898/2025.12.12.693809

**Authors:** Pravrutha Raman, Hana Khan, Janet M. Young, Rebecca Ferreira Alves, Maria Toro Moreno, Alice L. Li, Alice H. Berger, Toshio Tsukiyama, Harmit S. Malik

## Abstract

The Polycomb Repressive Complex 2 (PRC2) is an ancient, conserved chromatin-interacting complex that controls gene expression, facilitating differentiation and cellular identity during development. Its regulation is critical in most eukaryotes. EZHIP was recently characterized as a PRC2 inhibitor and ‘oncohistone mimic’ in mammals. Although *EZHIP* expression is typically restricted to the germline, its aberrant expression in pediatric brain tumors inhibits PRC2-mediated H3K27 methylation, driving disease progression. To gain a deeper understanding of its normal functions, we systematically examined *EZHIP* evolution across mammals using comparative genomics, synteny analyses, and motif discovery. Extending previous work, we find that *EZHIP* originated on the X chromosome and was retained there in most placental mammals, except in Afrotheria. In addition to the highly conserved H3K27M-like histone mimic motif, our motif analyses reveal six previously unidentified EZHIP motifs, including a putative nuclear localization signal, a serine-enriched region, and tandem repeats that are largely well-conserved in placental mammals. We hypothesize that these motifs are also critical to EZHIP’s functions, including in PRC2 interaction and inhibition. We show that *EZHIP* has evolved under strong diversifying selection in primates and underwent dynamic expansions and losses across species. Some paralogs, such as *EZHIP2* in primates, also evolved under positive selection. Based on its evolutionary attributes and germ-cell expression, we propose that *EZHIP* arose and evolved rapidly due to inter-parental conflict over fetal development *in utero* in placental mammals. Our work provides a foundation for further investigations into EZHIP’s multifaceted roles in mammalian reproduction and disease.

**Significance:** Histone modifications regulate genes to orchestrate cell differentiation and development. Their precise regulation is essential for normal development and growth. The Polycomb Repressive Complex 2 (PRC2) deposits the repressive H3K27 methylation mark, whereas EZHIP antagonizes PRC2 to de-repress gene silencing mediated by H3K27 methylation. While aberrant EZHIP activity in cancers is well documented, its normal germline functions, evolutionary origins, and divergence remain unclear. We show that EZHIP is a placental, X-linked innovation that diversified through positive selection and recurrent gene duplications, yielding germline-specific paralogs. We discover conserved motifs that may have critical roles in PRC2 control. Our findings suggest that parental control over *in utero* reproduction could have driven the origin and innovation of EZHIPs in placental mammals.

## Introduction

The Polycomb Repressive Complex 2 (PRC2) is a highly conserved multiprotein complex that plays a central role in gene silencing and regulating gene expression during development and differentiation (1, 2). The core subunits of PRC2 include EZH2 (<u>e</u>nhancer of <u>z</u>este <u>h</u>omolog <u>2</u>), the catalytic subunit responsible for methyltransferase activity, along with EED (<u>e</u>mbryonic <u>e</u>ctoderm <u>d</u>evelopment), SUZ12 (<u>su</u>ppressor of <u>z</u>este <u>12</u>), and RBBP4/7 (<u>R</u>etino<u>b</u>lastoma-<u>b</u>inding <u>P</u>rotein <u>4/7</u>, also known as RbAp46/48) (1, 2). Functionally, PRC2 is essential for establishing cellular memory and maintaining proper gene expression profiles during development by silencing genes involved in cell fate determination, including many developmental regulators and transcription factors (2). PRC2 catalyzes the methylation of histone H3 at lysine 27 (H3K27me3) (1). This epigenetic mark is associated with the formation of facultative heterochromatin and transcriptional repression (1). PRC2 has also been shown to silence transposable elements in some species (3, 4). In mammals, PRC2 plays critical roles in epigenetic processes, such as X chromosome inactivation, imprinting, and maintenance of stem cell identity (2). Dysregulation of PRC2, whether through loss or gain of function, has been linked to several diseases, including various cancers and developmental disorders, emphasizing the biological and clinical significance of this vital chromatin-modifying complex (2). Moreover, H3K27M ‘oncohistone’ mutations that render H3 impervious to PRC2 activity have been associated with disease progression in several cancers (5).

PRC2 activity is regulated by both internal feedback mechanisms and a variety of chromatin-associated factors, including post-translationally modified histones, noncoding RNAs, and accessory proteins, which help target PRC2 to specific locations or modify its catalytic activity (2, 6). These include histone modifications, such as H3K27me3 itself and H2AK119ub, as well as PRC2’s interactions with RNAs and nucleosome density, which control the timing and placement of gene silencing (2, 6, 7). Another mode of PRC2 regulation involves proteins that mimic unmodified histones to control the complex’s activity. For instance, PRC2 methylates accessory proteins JARID2 and PALI1 at specific lysine residues; their structural resemblance to the H3K27me3 mark enables them to bind EED and allosterically activate PRC2’s enzymatic action (6). Other proteins, such as AEBP2, also interact with PRC2 via histone tail-like motifs, further refining chromatin association and gene repression (6) .

Another ‘histone mimic’ that binds PRC2 is encoded by *EZHIP* (Enhancer of Zeste Homologs Inhibitory Protein; initially identified as *CXorf67*; also called *CATACOMB*, for catalytic antagonist of Polycomb) (8–12). Rather than activating PRC2, EZHIP acts as a competitive inhibitor of the PRC2 complex by interfering with EZH2 activity. In mouse embryonic stem cells, aberrant EZHIP expression constrains the spread of H3K27me3 in the absence of 5-methylcytosine DNA methylation, which can typically antagonize PRC2-dependent H3K27me3 (13). *In vitro* and *in vivo* studies showed that the C-terminus of EZHIP is both necessary and sufficient to repress PRC2’s function. This domain includes a conserved 14-amino-acid motif, called the K27M-like peptide (KLP), which contains the four amino acids VRMR that mimic an unmodifiable H3K27M motif (8, 9, 11). Mutating M (methionine) to a modifiable K (lysine) within the KLP restores wild-type H3K27 methylation levels, suggesting that this EZHIP motif is critical for inhibiting H3K27 methylation, similar to the H3K27M oncohistone mutation, which PRC2 can no longer modify (9, 11). The deletion of *EZHIP* increases global H3K27me2/3 methylation and is correlated with widespread changes in gene expression in cell lines (11, 12). Conversely, *EZHIP* overexpression lowers H3K27 methylation levels without affecting PRC2 expression, confirming that EZHIP affects PRC2 activity rather than its expression. EZHIP also appears to have minimal impact on PRC2 binding to chromatin (12) or its initial H3K27 methylation activity. Instead, it suppresses the spreading of PRC2 and H3K27me3 marks across the genome (9).

Abnormal activation of EZHIP in Posterior fossa type A (PFA) ependymomas, some osteosarcomas, diffuse midline gliomas, and rare non-CNS tumors is strongly implicated in oncogenesis due to global loss of the repressive histone mark H3K27me3 and epigenetic deregulation of PRC2 targets, even in the absence of the H3K27M oncohistone mutation (8, 14–16). A recent study using a *Drosophila* model showed that human EZHIP is an even more potent PRC2 inhibitor than the H3K27M oncohistone (17), suggesting that the EZHIP KLP may have higher affinity for PRC2 or that EZHIP has additional sequence features that enhance its inhibitory function.

Previous studies have primarily focused on the consequences of EZHIP’s aberrant expression in cancer. However, *EZHIP’s* normal expression is primarily restricted to oocytes and (to a lesser extent) testes, with little to no detectable expression in somatic tissues (12). Deletion of the mouse *Ezhip* results in a global increase in H3K27me2/3 during both spermatogenesis and late-stage oocyte maturation but does not cause gross infertility (12). Male knockout mice show a very mild defect in sperm motility. In contrast, female knockout mice show a progressive decline in fertility, with older females having smaller litter sizes and less healthy pups, presumably due to excessive accumulation of H3K27me3 (12). *EZHIP’s* biological function remains poorly studied. However, given its role in PRC2 regulation, *EZHIP’s* primary function may be to balance the germline epigenome between silencing and gene activation, thereby preserving totipotency and accurately transmitting epigenetic information to the next generation.

Unlike most PRC2 core proteins and cofactors, which are ancient and conserved across most metazoan genomes, EZHIP appears to be mammal-specific (18). Furthermore, EZHIP orthologs were reported to show no conservation beyond a 14-amino-acid KLP motif (9, 11). Yet functional experiments suggest that residues outside the KLP motif are important for EZHIP function. For example, full-length EZHIP restored PRC2 inhibition activity at lower concentrations than the KLP motif alone (9, 12). Moreover, EZHIP proteins lacking the KLP motif can still interact with EZH2 (11). Lastly, EZHIP’s function in DNA damage response in cancer is independent of the KLP motif (19). Despite these findings, *EZHIP’s* functional role in mammals remains unclear.

*EZHIP’s* absence outside placental mammals, its rapid sequence divergence except for the KLP motif, and its germline-restricted expression patterns all imply strong, lineage-specific evolutionary pressures. To gain insights into these pressures, we trace the origins and evolutionary dynamics of *EZHIP* across mammalian genomes. Consistent with previous results, we find that *EZHIP* arose and has been strictly retained on the X chromosome in most placental mammals, except in Afrotheria. Using motif analyses, we demonstrate that, in addition to the previously identified KLP motif, EZHIP encodes six well-conserved motifs and tandemly repeated sequences that vary in copy number and sequence across different *EZHIP* orthologs. We show that *EZHIP* evolves under diversifying selection in primates. We identify recurrent duplications of *EZHIP* in most mammalian lineages, resulting in *EZHIP* paralogs that have been retained for varying periods. For example, the mouse genome harbors up to 15 young X-linked *EZHIP* paralogs, while many carnivore and simian primate *species* have retained two independently duplicated copies of *EZHIP*. We identify a primate-specific autosomal paralog, *EZHIP2*, which evolves under diversifying selection like ancestral *EZHIP*. Like ancestral EZHIP, primate EZHIP2 can inhibit H3K27me3 modifications. However, unlike ancestral *EZHIP*, which is primarily expressed in oocytes, the primate paralog *EZHIP2* is expressed exclusively in testes. Our evolutionary analyses suggest a model in which *EZHIP* arose and diversified through sequence divergence and gene duplication, driven by competition between paternal and maternal genomes to gain control over *in utero* reproduction in placental mammals.

## Results

### *EZHIP* arose and has been retained in placental mammals

The single-exon *EZHIP* gene is located on the X chromosome in the human genome and encodes a 503-amino-acid EZHIP protein (Fig. 1A). The mouse X chromosome also encodes an *EZHIP* ortholog that encodes a 589-amino-acid protein. Previous studies have concluded that these two orthologs share only limited sequence similarity, with the histone mimic-like KLP motif as the only common feature identified so far. To systematically identify *EZHIP* homologs in mammals, we interrogated genome assemblies from a broad range of placental mammals, as well as a marsupial (opossum) and a monotreme (platypus). We used the human EZHIP sequence as a query to perform homology-based searches against each genome (see Methods). This approach successfully identified *EZHIP* homologs in most but not all placental mammals. Because of substantial divergence from human EZHIP, we could not identify *EZHIP* homologs in some placental mammal species, particularly rodents, despite the mouse genome previously being shown to encode *EZHIP*. Therefore, we performed iterative homology searches using EZHIP sequences from more closely related species as queries; this strategy enabled us to identify homologs in most queried placental mammals (Table S1). However, we could not identify *EZHIP* in marsupials or monotremes.

**Figure 1.**
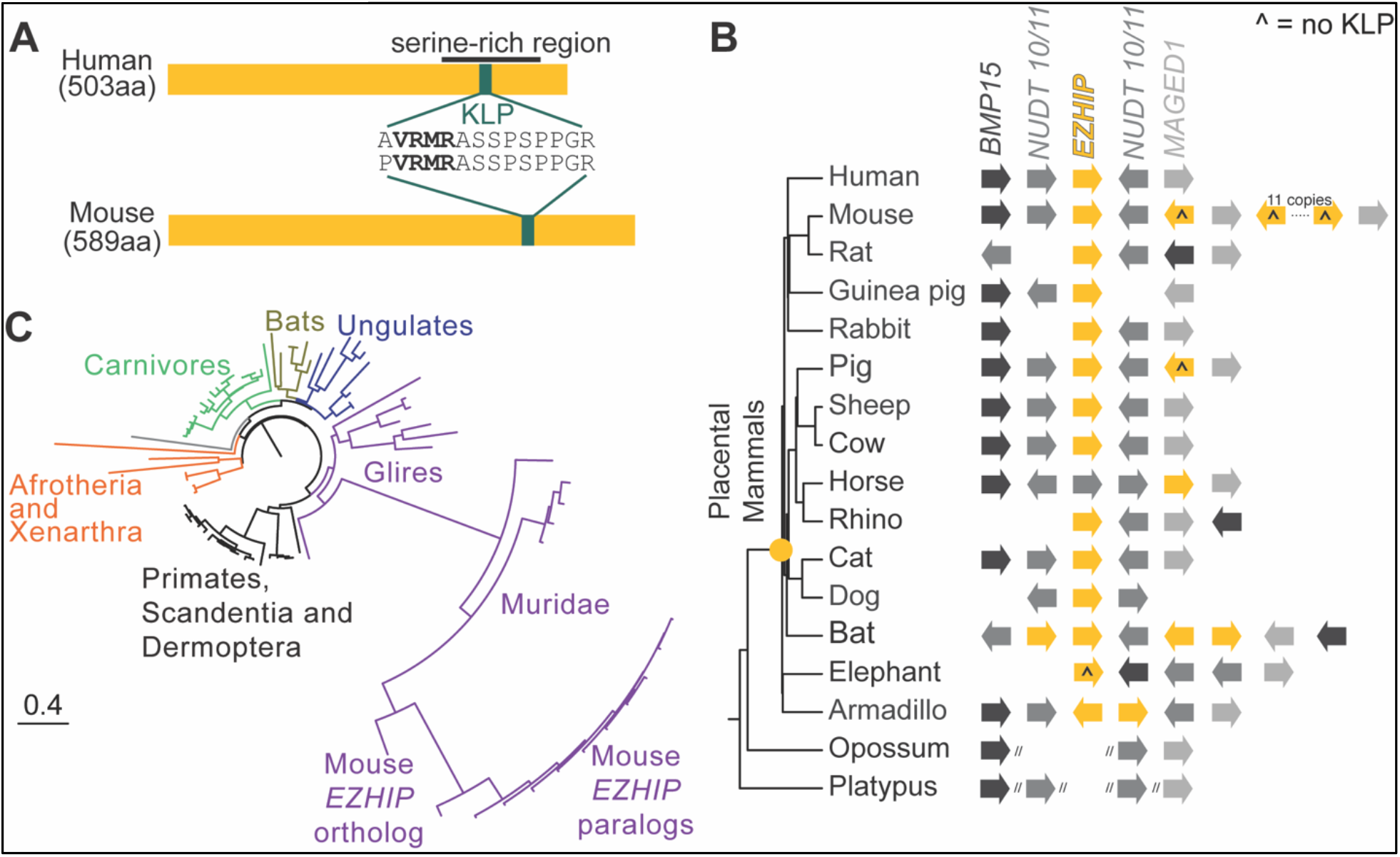
*Ezhip* arose in the last common ancestor of placental mammals. **A.** Schematics of human and mouse EZHIP proteins highlight previously identified domains (8, 9, 11, 12): a serine-rich region (black line) and a highly conserved 14-residue H3<u>K</u>27M-like <u>p</u>eptide (KLP) (green, with sequence shown) that contains the oncohistone-like VRMR (bold) motif. **B.** Shared syntenic locations of *EZHIP* genes (yellow) along with neighboring genes *BMP15, NUDT10/11,* and *MAGED1* (grey shades) in different mammalian species, shown next to an accepted mammalian phylogeny (22). *EZHIP* genes lacking KLP are denoted with a caret (^) symbol. A yellow dot on the species tree indicates the inferred origin of *EZHIP* in the common ancestor of placental mammals. We are unable to fully describe the syntenic locations in opossum or platypus due to short genomic scaffolds (two slashes). **C.** A maximum-likelihood protein phylogeny of syntenic *EZHIP* copies with complete ORFs across 64 representative mammalian species is presented as a circular phylogram, with different mammalian lineages highlighted: primates/ Scandentia (tree shrews)/ Dermoptera (flying lemurs), *Glires* (including Rodents), carnivores, ungulates, Chiroptera (bats), Afrotheria (incl. elephants), and Xenarthra (sloths, armadillos). The bottom-left scale bar below the phylogeny represents 0.4 substitutions per site. More details about the phylogeny, including individual taxon names, are shown in Supplementary Figure S4.

To confirm putative *EZHIP* orthologs, we relied on analyses of shared synteny (*i.e.,* conserved genomic neighborhood) in representative species. We used genes flanking *EZHIP* on the X chromosome, which are deeply conserved across mammals, to identify the syntenic ancestral locus. We identified the *EZHIP* gene at this shared syntenic location in most representative placental mammals from different lineages (Fig. 1B). This allowed us to conclude that *EZHIP* has been conserved at the shared syntenic location on the X chromosome of placental mammals. Furthermore, most placental mammals encode an intact *EZHIP* gene at the shared syntenic location, with an intact KLP motif.

However, Afrotherian species are a significant exception. Five species, including two tenrec species and two golden mole species, appear to encode an *EZHIP* gene with a KLP motif; dN/dS analyses indicate that the *EZHIP* gene in these species remains under purifying selection (Table S2, Data S1). Surprisingly, most other Afrotherian species either do not encode an intact *EZHIP* gene (*e.g.,* manatee, aardvark, elephant shrew, Fig. S1) or encode an *EZHIP* gene that lacks an intact KLP motif (*e.g.,* elephants, Fig. S2), indicated by a caret symbol (Fig. 1B, Fig. S1). We searched the entire elephant genome for sequences matching the KLP motif but found none. Thus, unlike other placental mammal lineages, many Afrotherian species do not appear to encode an intact *EZHIP* gene, nor any other gene with a KLP motif.

Unlike in placental mammals, we could not fully identify the *EZHIP* shared syntenic location in platypus or opossum genomes. In the platypus genome, we identified the flanking genes on completely distinct scaffolds because the genome assembly was incomplete. In the opossum genome, we found the flanking genes on two chromosomes: *NUDT10/11* and *MAGED1* were on chromosome X, while *BMP15* was on chromosome 8. Although we could identify the flanking genes, we did not find *EZHIP* in either genome. We also queried the KLP motif against these genomes but still could not identify this well-conserved EZHIP fragment. Based on these findings, we confirm that *EZHIP* arose in the last common ancestor of placental mammals and was strictly retained in placental mammals except *Afrotheria,* where it was differentially retained.

Most mammalian *EZHIP* orthologs have a well-conserved start codon. However, a multiple alignment unexpectedly revealed several cases of recurrent start-codon loss corresponding to the annotated human *EZHIP* start codon (Fig. S3). For example, *EZHIP* orthologs from gorilla, *Cercopithecinae* (Old World monkeys), and *Platyrrhini* (New World monkeys) have a mutation at the position corresponding to the annotated human *EZHIP* start codon (Fig. S3A). We hypothesize that monkeys and apes instead use a different methionine, which is five amino acids downstream of the annotated start and is well-conserved across primates. Dolphin *EZHIP* also has a mutation of the annotated start codon with a potential alternate start 17 codons downstream. Furthermore, several other *EZHIP* orthologs (tenrec, cat, and cow) encode proteins with divergent N-terminal sequences, and their start codons cannot be readily aligned with those of other placental mammals (Fig. S3B) due to frameshifts, deletions, or mismatches. Based on these findings, we infer that some placental mammal *EZHIP* orthologs have evolved alternate start codons.

Using a multiple alignment of EZHIP sequences from the syntenic region, we conducted phylogenetic analysis with PhyML (20, 21) (Fig. 1C, Fig. S4, Data S2). We found that their phylogenetic relationships matched our expectations based on the mammalian phylogeny (22), with very long branch lengths in *Glires*, particularly rodents. Our analyses thus confirm previous conclusions that *EZHIP* originated on the X chromosome in the last common ancestor of placental mammals (12). Although we find novel instances of loss in several Afrotherian species, *EZHIP* is otherwise retained in placental mammals, suggesting it may play important roles in their reproduction or development.

### *EZHIP* orthologs encode seven well-conserved motifs and independently derived tandem repeats

Previous studies have primarily focused on human and mouse EZHIP orthologs, which were thought to share only the 14-amino-acid KLP motif that includes the H3K27M-like VRMR sequence (Fig. 1A). Subsequent studies have noted the presence of other protein domains, such as a serine-rich region (10) or repetitive sequences (11), but these were considered idiosyncratic occurrences; most of the ∼500-amino-acid EZHIP protein remained uncharacterized.

We leveraged our comprehensive identification of EZHIP orthologs across placental mammals to uncover motifs shared by most orthologs to investigate the functional constraints that have shaped EZHIP function and divergence across mammals. Rather than relying on multiple alignments, whose accuracy could be affected by high sequence divergence or length differences, we instead relied on *de novo* identification of conserved protein motifs across mammals using the MEME suite (23), as previously described (24, 25) (see Methods). We identified 10 conserved motifs with high statistical significance across the majority of EZHIP orthologs in placental mammals (Fig. 2A). The newly identified motifs include residues that are strongly conserved across mammalian species (asterisks in Fig. 2A and Data S3).

**Figure 2.**
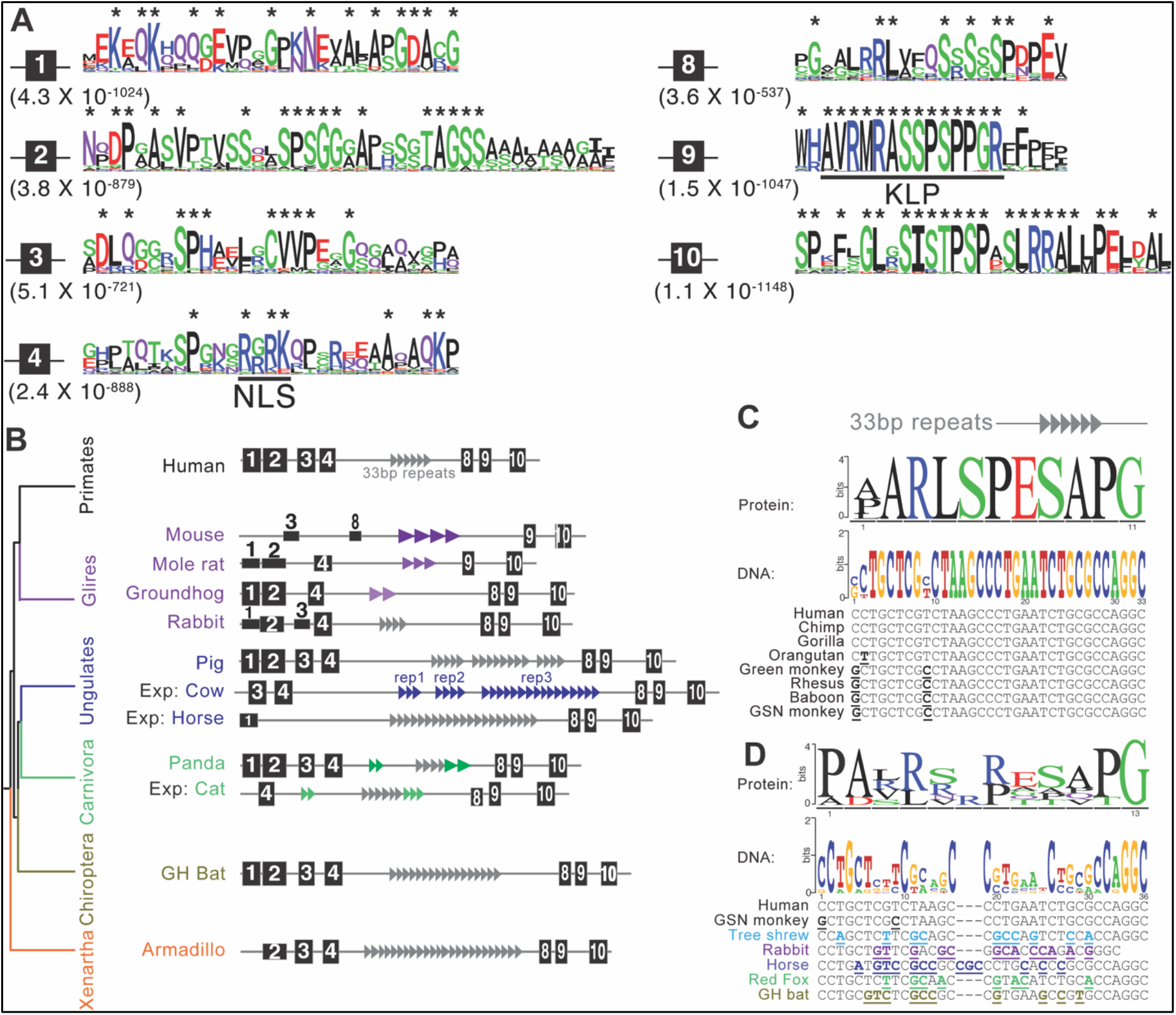
Conserved protein motifs and tandem repeats in EZHIP. **A.** Protein logo plots and associated e-values of eight motifs in *EZHIP* orthologs discovered using MEME analyses (23). Residue colors indicate their biochemical properties: hydrophobic (black), negatively charged (blue), positively charged (red), and polar (green), and residue heights correspond to their conservation (asterisks indicate residues conserved in >80% of motif occurrences found in mammalian species analyzed). Motif 4 includes a predicted <u>N</u>uclear <u>L</u>ocalization <u>S</u>ignal (NLS) from PSORT analyses (26), while motif 9 includes the H3<u>K</u>27-<u>L</u>ike <u>M</u>otif (KLP). **B.** A simplified species tree (22) of representative mammalian species groups (with the same color scheme as Fig. 1C) alongside selected EZHIP protein schematics (black filled boxes with numbers; motifs are drawn to scale). “GH bat” = greater horseshoe bat and “Exp:” = Exceptions within lineages. Supplementary Figure S5 shows schematics of all analyzed mammalian EZHIP proteins. Motif heights correlate with the significance of E-values, with shorter filled boxes representing weaker matches and partial motif matches indicated by dashed lines (*e.g.,* motif 10 in mouse that has small deletions). Since motifs 5-7 were repeated (Supplementary Figure S5 and S8), we used Tandem Repeat Finder (TRF) to detect repeats in *EZHIP*. Most *EZHIP* orthologs had varying numbers of an ancient, common 33-bp tandem repeat (gray triangles, also highlighted in logo plots in panel C and D), or lineage-specific repeats (green, blue, or purple). In some cases, such as cow, 3 different repeat types can be found. **C.** Nucleotide alignment of a single repeat unit across simian primates, along with logo plots showing nucleotide and amino acid conservation. Bold, underlined residues differ from the human sequence. **D.** Nucleotide alignment of a single repeat unit across a wider panel of mammals, along with logo plots showing nucleotide and amino acid conservation. (GSN monkey = golden snub-nosed monkey). Hyphens indicate gaps in the alignment. Residues that differ from the human sequence are bold, underlined and colored (to represent lineage). Also see Supplemental Figure S9.

We mapped the occurrence of the motifs onto each of the orthologs using the MAST software within the MEME suite (Fig. 2B, Fig. S5). We numbered these motifs sequentially from N- to C-termini. Based on this numbering scheme, the previously identified KLP falls within motif 9. As expected, this motif had the highest statistical significance of all ten motifs; it is present and nearly identical across all orthologs, except in elephant EZHIP, where we previously found it to be lost (Fig. 1B, Fig. S4). The MAST analysis revealed that, in addition to KLP (within motif 9), several additional motifs are broadly conserved across most mammalian EZHIP orthologs (Fig. 2B, Fig. S5). Using PSORT analyses (26), we also identified a putative nuclear localization signal (NLS) within motif 4 (Fig. 2A). We used AlphaFold to understand where motifs lie in the predicted structure of EZHIP (Fig. S6). We found that EZHIP is highly unstructured except for low-confidence structured regions within motif 1 and motif 9. However, given the extremely low confidence in this predicted EZHIP structure, we cannot generate any testable hypotheses from it.

Among the ten motifs, MAST analysis revealed that three of the motifs (5–7) were present more than once in each ortholog (Fig. S5). In contrast, the other seven identified motifs (motifs 1-4, 8-10) are represented only once in most EZHIP orthologs. These seven motifs include amino acid residues that are highly conserved across mammals (asterisks in Fig. 2A indicate the same amino acid is found in >80% of motif occurrences). However, there were some striking exceptions to the near-universal conservation of these seven EZHIP motifs, most notably among *Glires* (including rodents) (Fig. 2B, Fig. S5). Most EZHIP orthologs in *Glires* retain motifs 4, 8, 9, and 10, but with much lower statistical support than in other mammals. Motifs 1, 2, and 3, which were otherwise prevalent across other mammalian EZHIP orthologs, are much less conserved in *Glires* and sometimes absent in *Muridae*. This low retention of otherwise well-conserved motifs in *Muridae* likely reflects high EZHIP divergence within this lineage, as shown by long branch lengths for orthologs in the phylogenetic analyses (Fig. 1C, Fig. S4). Furthermore, we were unable to identify robust rodent-specific motifs when we limited our analyses to rodent EZHIP orthologs. *Glires*, particularly *Muridae*, are the most striking examples of EZHIP motif loss, but such motif loss also occurs idiosyncratically in other lineages (“Exp:” in Fig. 2B indicate exceptions, Fig. S5). Although it remains unclear how the loss of these motifs alters EZHIP functions, we observed that N-terminal motifs are more likely to be lost than C-terminal motifs. Previous studies showed that the C-terminal EZHIP domain encompassing the KLP motif is necessary and can be sufficient for EZHIP’s PRC2 inhibitory function (8, 11), suggesting that even when some N-terminal motifs are lost, EZHIP’s PRC2 inhibitory function is likely retained.

Previous work has suggested a serine-rich region surrounding the KLP in human EZHIP (10). Consistent with this prediction, we observed high conservation of serine residues in motifs 8 through 10 and observed an increased frequency of serine residues after the KLP in EZHIP orthologs (some representative orthologs shown in Fig. S7). This pattern of serine enrichment at the C-terminus is also observed in elephant EZHIP, which has lost the KLP motif. These data suggest that a serine-rich region may be crucial for EZHIP function.

Our finding that motifs 5 to 7 are repetitive within most orthologs (Fig. S5, Fig. S8) suggested they might be part of a repetitive region in EZHIP. Indeed, Tandem Repeat Finder (TRF) (27) analyses identified tandem repeat sequences present in EZHIP sequences from ∼50 mammalian species. We manually curated the results of this analysis to determine the smallest repeat unit to map the number of repeats in representative EZHIP orthologs (Fig. 2B). Repeat unit copy numbers vary dramatically across EZHIP orthologs, ranging from as few as 3 copies in some bats and carnivores to as many as 21 copies in some ungulates. Using the Genome Aggregation Database (gnomAD) (28), we find that the number of EZHIP tandem repeats can vary even within the human population. For instance, although the most common EZHIP allele in humans has 6 repeats, other alleles only have 5 repeats.

Using iterative sequence alignments within a lineage (*e.g.,* among carnivores) and between lineages (*e.g.,* between carnivores and primates), we identified patterns of common and lineage-specific repeats. Based on these comparisons, we infer that most EZHIP orthologs contain a related ∼33 bp repeat, suggesting this was a feature of EZHIP since its origin. For example, we can identify related tandem repeats in primates, ungulates, some carnivores (such as *Canidae*), and bats (gray triangles, Fig. 2B). The repeat unit is relatively well conserved at the nucleotide and protein levels, both within a species (Fig. S9) and across lineages, with >95% pairwise nucleotide identity within simian primates (Fig. 2C) and >64% pairwise nucleotide identity between lineages (Fig. 2D). In addition to the ∼33 bp repeat present across most mammalian EZHIP proteins, we found that some lineages have also evolved ‘new’ lineage- or even species-specific tandem repeats (represented in lineage-specific colors, Fig. 2B), which show sequence, length, and copy-number divergence. For example, although the ancestral 33 bp repeat is present in *Canidae*, other carnivores have lost it and gained three new repeats (33 bp, 39 bp, and 66 bp). Similarly, while most ungulates retained the ancestral 33-bp repeat, species such as *Bos taurus* (cow) lost it and gained three new repeat families, each 36 bp long (Fig. 2B). Lastly, although rabbit EZHIP encodes the ancestral 33-bp repeat, it is absent in rodents, which gained new repeats, each differing in length and sequence (Fig. 2B). Given the dynamics of tandem repeats and the long evolutionary timescales, we cannot rule out the possibility that these new repeats are themselves ancestrally derived from the original 33 bp repeat. Based on these analyses, we conclude that tandem repeats appear to have been a feature of EZHIP orthologs throughout placental mammal evolution, along with the six new motifs we identified.

We also leveraged these protein motifs to perform more sensitive searches of selected genomes and predicted proteomes to identify additional remote homologs of all or part of the EZHIP gene. We used three methods (Hidden Markov Models, FIMO, and MAST searches in the MEME suite; see Methods) but found no convincing homology in species outside of placental mammals, confirming our inferred age of EZHIP. Other than EZHIP and some recent duplicates identified using BLAST searches (see below), we found no evidence for additional paralogs in selected placental mammal genomes. We also looked for additional genes containing only the KLP motif and found no clear matches, including in the genomes of Afrotherian species where EZHIP lacks the KLP. The evolutionary origin of EZHIP and the KLP motif therefore remains unclear.

### *EZHIP* has evolved under diversifying selection

Given its retention, we infer that *EZHIP* performs a vital function in the germline tissues of all placental mammals. However, our analyses also revealed an apparent disparity among lineages in EZHIP evolution, with some lineages exhibiting long branches (Fig. 1C, Fig. S4) or lacking ancestral protein motifs (Fig. 2, Fig. S5). To investigate this disparity further, we compared the rate of EZHIP protein divergence across different mammalian lineages, including primates, rodents, carnivores, and ungulates, spanning nearly 70 million years of evolution in each lineage (Fig. 3A, Table S3). For our analyses, we measured the pairwise identity of EZHIP orthologs relative to a ‘reference’ sequence within that clade. For example, we compared the pairwise identities of various primate EZHIP orthologs to human EZHIP. We then compared this protein divergence to the estimated species divergence times obtained from TimeTree (29). To be conservative and consistent, we chose the most conserved ortholog when multiple paralogs were found in the shared syntenic location. We carried out similar comparisons in rodents (relative to mouse EZHIP), ungulates (relative to goat EZHIP), and carnivores (relative to dog EZHIP). Our analyses show that EZHIP orthologs in primates, ungulates, and carnivores diverge rapidly, with as little as ∼60% conservation among orthologs, in stark contrast to EZH2, the PRC2 component that interacts with EZHIP, which is >90% identical across primates. Rodent EZHIP orthologs show even faster divergence, with only 40% identity over 20 MY (Fig. 3A), whereas rodent EZH2 remains highly conserved, with >98% identity across the same species. Our results are consistent with both our EZHIP phylogeny and motif analyses (Fig. 1C, Fig. 2B, Fig. S4) and with previous evolutionary analyses highlighting the much faster rate of protein divergence in rodent genomes (30).

**Fig. 3.**
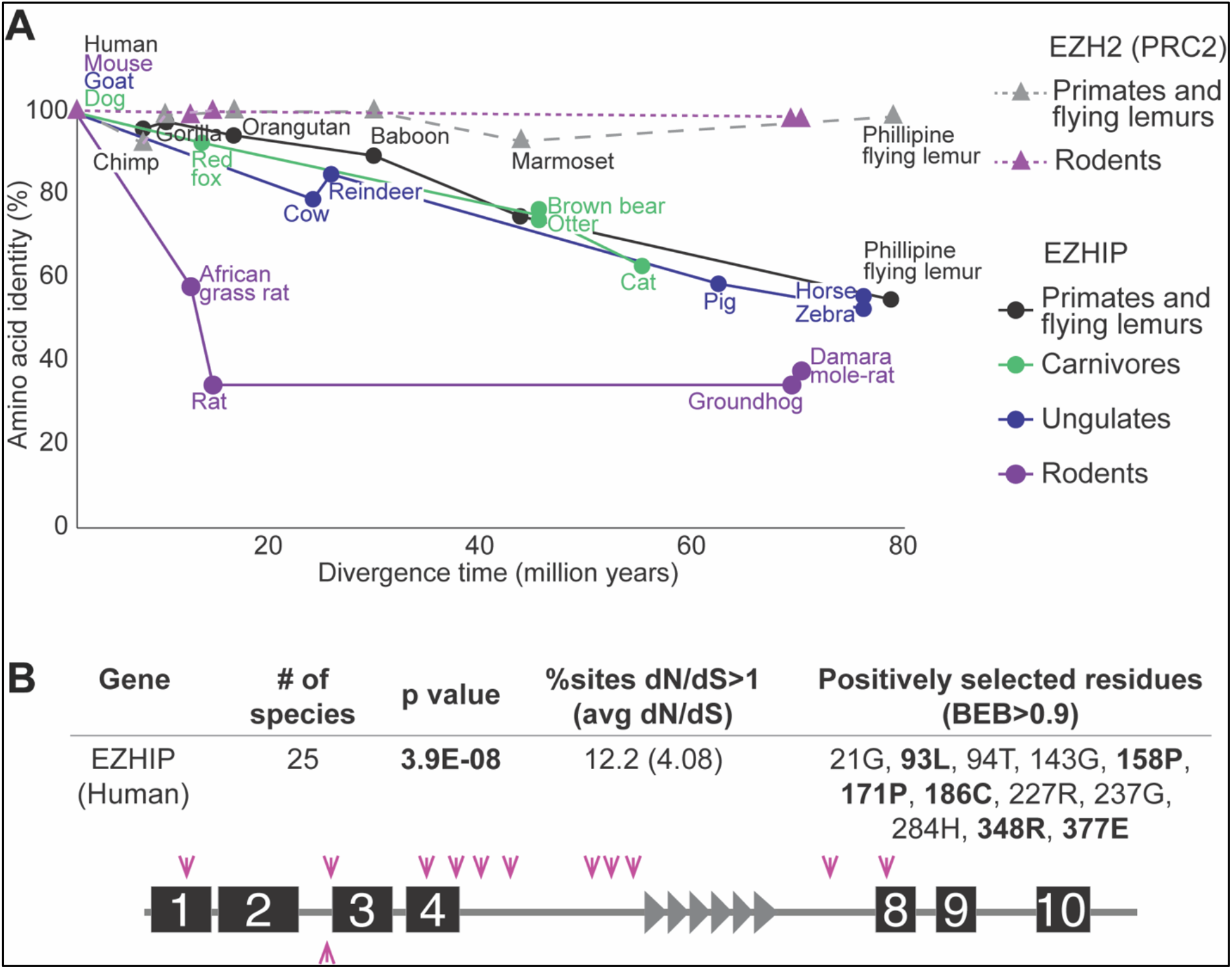
Rapid evolution of EZHIP. **A.** Species divergence time estimated from TimeTree (29) (in millions of years) on the X-axis is plotted against pairwise percent amino acid identity of EZHIP orthologs in primates, rodents, ungulates, and carnivores relative to human, mouse, goat, and dog, respectively (circles, colors match Fig. 1C). A similar divergence plot is shown for EZH2, a component of the PRC2 complex, in primates (gray triangles with dashed line) and rodents (purple triangles with dotted line). **B.** Individual residues in EZHIP identified as evolving under diversifying selection in simian primates, according to PAML (BEB>0.9 (31)). Residues that were also identified by FUBAR (33) are shown in bold. P-value showing a better fit to PAML Model 8 over Model 8a for the entire *EZHIP* gene, the percentage of codons estimated to evolve with dN/dS>1, and their average dN/dS are indicated. A schematic of the human EZHIP protein, highlighting the locations of residues evolving under diversifying selection, is shown below. Fig. S10 and Table S4 provide additional details of these analyses.

High EZHIP divergence across mammals could indicate a loss of constraint, although this is unlikely given *EZHIP’s* high retention in placental mammals. We examined selective constraints acting on *EZHIP* by comparing the normalized ratio of amino acid-altering changes (non-synonymous, dN) to amino acid-preserving changes (synonymous, dS). Neutrally evolving sequences are expected to have dN/dS close to 1, whereas sequences under selective constraint (purifying selection) are expected to have dN/dS < 1. Using a set of mammalian species from the lineages selected above (Fig. 3A), we compared the relative likelihoods of codon models assuming neutral versus non-neutral evolution. Specifically, we used PAML’s codeml program (31) to estimate the likelihood of a simple evolutionary model (model 0), in which all sequences and codons are assumed to have the same dN/dS ratio. We compared model 0, with the likelihood of dN/dS fixed at 1 (neutral, as expected for a pseudogene), to that of dN/dS estimated from the alignment (∼0.85). This test revealed evidence of weak purifying selection on *EZHIP* across mammals (Table S2, Data S4).

Despite this overall signature of purifying selection, a subset of *EZHIP* codons could evolve under positive selection (dN/dS >1). To investigate this possibility, we analyzed sequences from simian primates, which have an ideal level of evolutionary divergence for codon-by-codon analyses (32). We analyzed intact EZHIP sequences from 25 simian primates, excluding repetitive regions, as they are difficult to align confidently (Table S4, Data S5). We assessed codons undergoing positive selection using maximum-likelihood analyses with PAML (31) and FUBAR (HyPhy package (33, 34)). PAML analyses revealed that 12.2% of EZHIP sites evolve with an average dN/dS of 4.08 (Fig. 3B). A subset of these sites met the Bayes Empirical Bayes (BEB) statistical threshold of 0.9. FUBAR analyses identified nearly the same sites, and some additional sites under positive selection (Fig. 3B, Table S4, Fig. S10). About half of the positively selected residues in EZHIP fall within the protein motifs we identified earlier in this study, ruling out the possibility that primate *EZHIP’s* positive selection is an artifact of poor alignments. Instead, our findings show that *EZHIP* is undergoing purifying selection at the whole-gene level, but that a subset of sites has been subject to diversifying (positive) selection, at least in primates.

### *EZHIP* has been recurrently duplicated in placental mammals

Previous studies have focused exclusively on single-copy *EZHIP* orthologs located in the same syntenic region. However, our analyses also identified multiple within-species duplications of *EZHIP*, creating paralogous copies (Fig. S11). Using protein sequence alignments of all identified *EZHIP* orthologs and paralogs (Table S1), we manually curated all paralogs as being either intact, incomplete due to gaps in genome assembly (‘*i*’, Fig. S11), pseudogenes with obvious disruptions in the open-reading frame (boxes containing crosses, Fig. S11), or intact but lacking a KLP motif (caret, Fig. S11). We note that incomplete *EZHIP* paralogs or those lacking a KLP motif may still be selectively retained if they encode functional proteins, although an intact KLP motif has previously been shown to be essential for PRC2 regulation (9, 11).

We found more than one *EZHIP* homolog at the ancestral shared syntenic location on the X chromosome in multiple species: Philippine flying lemur, African grass rat, greater horseshoe bat, rhinoceros, armadillo, and sloth (Fig. S11). Other *EZHIP* paralogs are found at distinct genomic locations: on sex chromosomes (blue boxes), on autosomes (pink boxes), or in unmapped regions (black boxes). The most spectacular example of *Ezhip* duplications is seen in the mouse genome, which encodes 15 X-chromosomal *Ezhip* paralogs (Fig. 1B, Fig. S11, Table S1). Four of these mouse *Ezhip* paralogs appear to be pseudogenes lacking an intact open reading frame, while the remaining 11 mouse *Ezhip* paralogs appear to be intact but lack a KLP motif (Fig. 1B, caret symbols; Fig. S12).

Phylogenetic analysis of EZHIP and its paralogs using maximum-likelihood methods (Fig. S13, Data S7) revealed that most paralogs are species-specific; they are more closely related to the *EZHIP* orthologs at the ancestral syntenic location in the same genome than to *EZHIP* genes from other species. However, not all *EZHIP* duplication events are young. We identified at least two instances in which *EZHIP* paralogs have been selectively retained for long periods. The first of these is in carnivores, where an *EZHIP* paralog appears to have been retained for 60 million years (Fig. S14). We traced this *EZHIP* paralog to a single X-chromosomal duplication event in the last common ancestor of *Caniformia* (Fig. S14), a sub-clade of carnivores. This *EZHIP* paralog has undergone pseudogenization in the dog genome and is absent from the cat genome, even at syntenic locations. It has undergone further duplication in *Ursidae*, *Mustelidae*, and *Pinnipedia,* resulting in additional intact or pseudogenized copies that cluster together in the phylogeny (Fig. S13). Most of these paralogs lack an intact KLP motif, suggesting they may have lost their PRC2-inhibitory functions.

A second ancient *EZHIP* duplication, which we named *EZHIP2,* arose once on an autosome (human chromosome 5) in the last common ancestor of *Haplorhini* (simian primates and tarsier) and appears to have been largely retained for ∼40 million years (Fig. 4A, Fig. S13). *EZHIP* and *EZHIP2* loci share no homology beyond the gene itself, suggesting that *EZHIP2* arose via a retrotransposition event. *EZHIP2* appears to be retained in simian primates except in two clear instances: bonobo/chimpanzee and marmoset/squirrel monkey, which lack open reading frames and KLP motifs. Otherwise, *EZHIP2* genes from Old World monkeys and a subset of hominoids encode motifs 3-4, with an intact NLS region, and 8-9, including a well-conserved KLP motif (Fig. 4A, Fig. S15A). To identify tandem repeats, we used TRF, as we did for *EZHIP*. With a slight modification to relax of our previously used TRF criterion (see methods), we could identify tandem repeats in *EZHIP2*, which have diverged from ancestral *EZHIP* (Fig. 4A, Fig. S15A). The status of human *EZHIP2* remains less certain. An ∼800bp region within the human *EZHIP2* ORF underwent an inversion, leaving a much shorter open reading frame (88 amino acids) that only encodes motifs 8 and 9, including an intact KLP motif (Fig. S15B). As a result, human *EZHIP2* now encodes a much shorter open reading frame, using a novel putative start codon upstream of motif 8.

**Figure 4.**
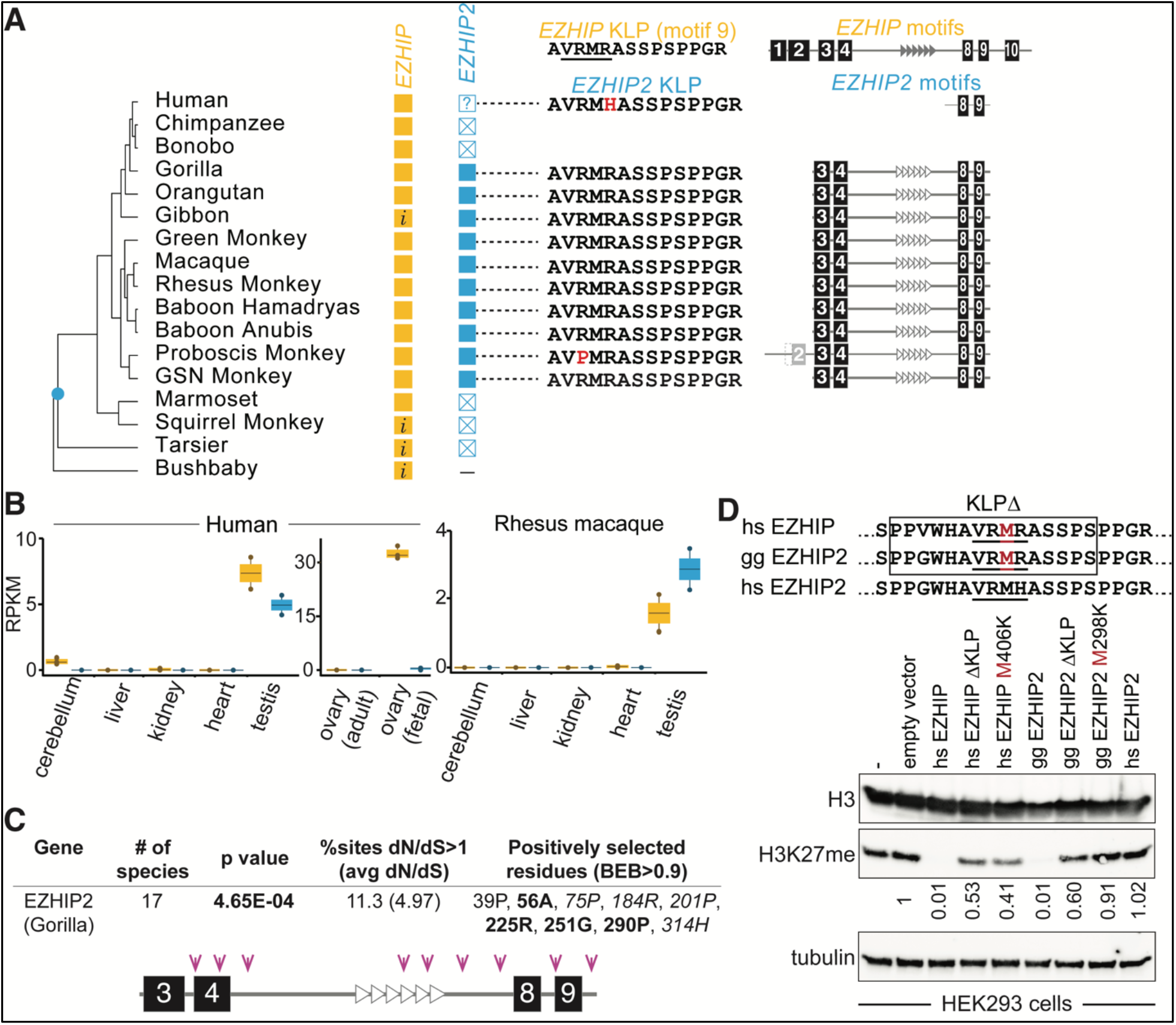
*EZHIP2* is a primate-specific *EZHIP* paralog. **A.** Left: retention of *EZHIP* (yellow boxes) or *EZHIP2* (blue boxes) is illustrated alongside a primate species tree (“X” = putative pseudogenes; “*i*” = incomplete sequences; “?” =truncated ORF). Since *EZHIP2* could not be identified in bushbaby through homology or synteny analyses, we infer that *EZHIP2* arose in Haplorhini (blue dot in species tree). Middle: aligned sequences show conservation of the KLP motif (red indicates residues that differ from the KLP in human EZHIP, top). Right: most primate EZHIP2 proteins appear to have retained a subset of the conserved protein motifs that are found in human EZHIP (top). Tandem repeats (empty triangles) can be identified in EZHIP2 using less stringent parameters than for EZHIP. See Supplementary Figure S15 for additional details. **B.** Analyses of publicly available RNA-seq datasets indicate that *EZHIP* (yellow) is expressed in fetal ovaries and testes, and slightly in the cerebellum, whereas *EZHIP2* (blue) appears to be exclusively expressed in testes. RPKM (reads per kilobase per million reads sequenced) is plotted for each tissue, with standard error bars. **C.** Individual residues in gorilla *EZHIP2* identified as evolving under diversifying selection in simian primates, according to PAML (BEB>0.9 (31)) or FUBAR (italic font (33)) or both (bold). Also indicated are the p-value showing a better fit to PAML Model 8 over Model 8a for the entire *EZHIP2* gene, the percentage of codons estimated to evolve with dN/dS>1, and their average dN/dS. A schematic of the gorilla EZHIP protein, highlighting the locations of residues evolving under diversifying selection, is shown below. Supplementary Figure S18 and Supplementary Table S4 provide additional details of these analyses. **D.** Western blot analyses of H3K27me3 modification, total H3, or tubulin from HEK293 cells expressing *Homo sapiens* EZHIP (hsEZHIP), *Gorilla gorilla* EZHIP2 (ggEZHIP2), or *Homo sapiens* EZHIP2 (hsEZHIP2) reveal that gorilla EZHIP2, like human EZHIP, can inhibit H3K27me3 modification. Deletion of KLP (top, boxed) or mutation of M (top, red font) to a K in the oncohistone-like VRMR motif results in loss of this inhibition activity. In contrast, human EZHIP2, which is significantly truncated, has no impact on H3K27me3 modification levels.

We analyzed *EZHIP* and *EZHIP2* expression using public RNA-seq data (Table S5) from human and rhesus macaque samples (Fig. 4B). As in previous work (12), we found that *EZHIP* and *EZHIP2* are almost undetectable in most somatic tissues. However, unlike prior work, we detected low levels of human *EZHIP* expression in the cerebellum, a tissue where *EZHIP* is often mutated or overexpressed in PFA ependymomas (8). Consistent with previous work, we find that human *EZHIP* expression is highest in reproductive tissues, both testes and ovaries, with higher expression in fetal ovaries than in adult ovaries (Fig. 4B) (12). In contrast, human *EZHIP2* expression appears to be limited to the testes, with no expression in the female germline or somatic tissues. This testicular *EZHIP2* expression pattern is also recapitulated in rhesus macaques, which encode a more intact *EZHIP2* (Fig. 4B). Both *EZHIP* and *EZHIP2* are expressed early in human spermatogenesis (Fig. S15C), with no expression detected in post-meiotic sperm. *EZHIP2* expression increases slowly, peaks in pre-meiotic and meiotic sperm, then decreases post-meiosis. Lastly, while human *EZHIP* is highly expressed in the embryo, we did not detect *EZHIP2* expression in the embryo (Fig. S15C). Together, these differential expression patterns suggest that *EZHIP* and *EZHIP2* may have evolved unique expression patterns and potentially, non-redundant functions in the male germline in most simian primates.

We also assessed *EZHIP* and *EZHIP2* expression levels in human cancer samples. *EZHIP* was first identified due to its elevated expression in PFA ependymomas (10). Since that discovery, *EZHIP* enrichment has been observed in several other cancers, including osteosarcomas, diffuse midline gliomas, non-CNS tumors, and small cell lung cancers (8, 14–16). We took advantage of the large number of cancer cell lines and samples sequenced by the Cancer Cell Line Encyclopedia (CCLE) (35) and The Cancer Genome Atlas (TCGA) (36) projects to investigate *EZHIP* and *EZHIP2* expression in a wide array of cancers. Although most cancer cell lines lack *EZHIP* expression (Fig. S16), our comprehensive analyses comparing TCGA tumor and normal tissues show that *EZHIP* expression is robustly detectable across a much wider array of cancer types than previously demonstrated (Fig. S17). In contrast, *EZHIP2* was barely detectable in these samples (Figs. S16 and S17).

Another approach to assessing whether *EZHIP2* is functional is to examine whether it evolves under selective constraints, as *EZHIP* does. We analyzed full-length *EZHIP2* sequences from 17 non-human simian primates, excluding all primates with obvious pseudogenizing mutations and the significantly truncated human *EZHIP2* (Table S4, Data S9). Our analyses allow us to reject the hypothesis that *EZHIP2* evolves neutrally. Using PAML analyses, we inferred that *EZHIP2* evolves with an overall dN/dS of 1.35 with a higher likelihood (p-value=0.02) than the null hypothesis of neutral evolution (dN/dS =1). Consistent with the whole-gene estimate of diversifying selection, our codeml analyses also found that 11.3% of *EZHIP2* codons evolve with a dN/dS of 4.97 (Table S4). Several sites meet a Bayes Empirical Bayes threshold of 0.9. FUBAR analyses identified some of these sites, along with additional sites that likely evolved under positive selection (Fig. 4C, Table S4, Fig. S18). Our selection analyses find evidence that *EZHIP* paralogs have undergone recurrent innovation in placental mammals, both through site-specific positive selection (in primates) and recurrent whole-gene duplications. Thus, although EZHIP and its paralogs are strictly retained in placental mammals, they show signatures of participation in biological processes that require constant adaptation.

Despite its truncated nature, we did not want to assume that human EZHIP2 is incapable of inhibiting PRC2. Indeed, previous experiments had shown that an intact KLP motif with some surrounding segments can be sufficient for H3K27me3 inhibition (9, 11). To test whether EZHIP2 is functionally capable of PRC2 inhibition (8, 9, 11, 12), we assessed whether its expression in human cells could reduce H3K27 methylation. For this, we expressed human *EZHIP*, gorilla *EZHIP2,* or human *EZHIP2* in HEK293T cells (Fig. 4D). Given the uncertain status of human *EZHIP2*, we chose gorilla *EZHIP2* as the most closely related intact ortholog (because chimpanzee and bonobo *EZHIP2* have been pseudogenized). Consistent with previous results (8, 9, 11, 12), expression of human EZHIP led to a marked loss of H3K27me3. This decrease depended on the KLP, specifically the VRMR motif, because deletion of the KLP (ΔKLP) or mutation of M to K (M406K) within the VRMR oncohistone-like motif restored H3K27me3 methylation (Fig. 4D). We found that this PRC2 inhibitory function is retained in gorilla EZHIP2. Expression of gorilla *EZHIP2* led to a marked decrease in the H3K27me3 mark, suggesting that it has retained PRC2 inhibitory functions. Like ancestral EZHIP, deletion of the KLP (ΔKLP) or mutation of M to K (M298K) within the VRMR motif of gorilla EZHIP2 restored H3K27me3 levels. Comparing gorilla EZHIP and EZHIP2 sequences (Fig. S15) provides further clues about motifs that are crucial for PRC2 inhibitory functions. Gorilla EZHIP2 has lost motifs 1, 2, and 10 (Fig. S15A) when compared to EZHIP, suggesting that these motifs are not necessary to inhibit PRC2 or H3K27 methylation. However, a majority of the strongly conserved EZHIP residues we identified in our motif analyses are still retained in EZHIP2 (asterisks in Fig. 2A and Fig. S15A), supporting our hypothesis that these residues may be crucial for PRC2 inhibitory functions. Unlike gorilla *EZHIP2*, human *EZHIP2* expression did not alter H3K27 methylation levels. These data suggest that *EZHIP2* was likely pseudogenized recently, at least with respect to its PRC2-inhibiting activity, in the common ancestor of humans, chimpanzees, and bonobos. In contrast, the H3K27-methylation inhibition assay and the signature of diversifying selection strongly suggest that *EZHIP2* is still functional in regulating PRC2 in most simian primates. Thus, *EZHIP* has undergone repeated innovation via both sequence evolution and gene duplication in placental mammals.

## Discussion

In this study, we revisited the exciting discovery of EZHIP, a ‘histone mimic’-based negative regulator of PRC2 (8, 9, 11, 12), by characterizing its evolution across more than 80 mammalian species to understand its functional significance. Our study reveals that *EZHIP* originated on the X chromosome in the last common ancestor of placental mammals (∼100 MYA) and has been retained in most placental mammals since its origin, except in several Afrotherian species. Beyond the previously identified KLP motif, we identified six additional conserved motifs and lineage-specific tandem repeats retained across most placental mammals. Most strikingly, *EZHIP* exhibits strong signatures of genetic innovation, undergoing both diversifying selection in primates and recurrent duplications and post-duplication losses across mammalian lineages. For example, *EZHIP* paralogs include 11 X-linked paralogs in mice and two independent autosomal duplications retained for long periods in carnivores and primates. Primate *EZHIP2* is most highly expressed in the testes, has evolved under positive selection, and retains the ability to inhibit PRC2. Although the strict retention of *EZHIP* in placental mammals suggests an important, conserved function, our discovery of constant innovation implies that *EZHIP* and its paralogs are also subject to changing selective pressures that require ongoing adaptation. One reason for EZHIP divergence across eutherian lineages could be differences in placentation patterns across eutherian orders (37), although current genetic knockout evidence in mice suggests EZHIP may not be important for placental development (38). Alternatively, recurrent signatures of innovation may suggest EZHIP’s involvement in genetic conflict and could provide vital clues about its function.

Our study has implications for several aspects of *EZHIP’s* multifaceted functions, starting with its protein architecture. Systematic truncations of EZHIP expressed in human cells had previously revealed the necessity but insufficiency of the KLP motif alone to inhibit H3K27 methylation (8, 9, 11) or to bind PRC2 (9, 11). Together, these data suggested that EZHIP must have additional regions that facilitate its sole well-characterized function: binding to and inhibiting PRC2. Yet, most previous studies had focused almost exclusively on the conservation of the KLP motif. Our study identifies several additional motifs, whose conservation suggests they could also contribute to EZHIP’s role in regulating PRC2. This may occur by increasing binding avidity to the PRC2 complex, altering its localization to different genomic loci, or repressing PRC2 at specific (imprinted) loci, potentially changing the landscape of H3K27me3 modifications across germline genomes. We posit that a focus on the highly divergent mouse EZHIP, one of the earliest EZHIP orthologs identified, may have obscured the identification of these conserved motifs. Our work highlights the power of *de novo* motif identification over alignment-based methods to identify regions of homology and functional constraint in highly divergent proteins such as EZHIP. We predict that mutating these well-conserved motifs may reveal key insights into EZHIP’s functions in future studies. We also identified tandem repeats that are prevalent across EZHIP. We speculate that these repeats may contribute to the allosteric regulation of KLP binding to PRC2 (or EZH2) with differential affinities to fine-tune H3K27me3 spreading across the genome. The differences in H3K27me3 distributions across species and germ cells (39–43) raise the intriguing possibility that such repeat-rich regions serve as an evolutionary playing field in which mutations can strengthen or weaken PRC2 interactions. Our findings also suggest that *EZHIP* orthologs or paralogs may have PRC2-independent functions that contribute to their evolutionary retention. For example, aberrant *EZHIP* expression in PFA ependymomas has recently been shown to suppress homologous recombination-mediated DNA repair, independent of the KLP motif, suggesting that EZHIP may modulate the DNA damage response (19).

Our findings have additional implications for EZHIP’s role in cancer. *EZHIP* was initially identified because of its mutation or misexpression in cancers, particularly posterior fossa type A (PFA) ependymomas (8, 10, 44). Our analyses reveal that *EZHIP* is expressed in an even wider array of cancers than previously believed (Fig. S17). Our identification of the *EZHIP2* duplicate, which encodes an intact KLP motif, raised the possibility that it may also be misexpressed in some cancers, where it might drive cancer progression despite encoding a truncated protein. Although human *EZHIP2* appears unable to repress PRC2 activity in cell culture, fusions between truncated EZHIP and transcription factors have been predicted to result in aberrant PRC2 localization in cancers (44). One such fusion identified in endometrial stromal sarcomas occurred between a truncated EZHIP (containing only motifs 8-10) and MBTD1, a chromatin reader of the NuA4 histone acetyltransferase complex (44). Our discovery of *EZHIP*2, which also encodes most of motifs 8-9, suggests that gene fusions involving human *EZHIP2* might still be able to alter PRC2 localization and function in cancers.

Our study also has implications for previous and future genetic studies of *EZHIP* function. The best-known system in which *EZHIP’s* canonical function has been elucidated is in mice (12), where *Ezhip*’s knockout phenotypes were modest, given the strict retention of *EZHIP* in placental mammals: no effect on male fertility and a modest age-dependent decline in female fertility. Our finding that the mouse genome encodes at least 11 additional intact *Ezhip* paralogs (lacking the KLP motif) suggests that single gene knockouts may be insufficient to study the consequences of a complete loss of *Ezhip* activity in mice. Previous studies have demonstrated differences in imprinting patterns between rodents and other eutherian mammals as well as differences in the persistence of gene repressive marks in the embryo after genome activation (45–47). One possibility is that these imprinting differences facilitated the accelerated evolution of EZHIP in rodents. Future genetic perturbations to study *Ezhip* might have to account for all mouse *Ezhip* homologs, or use an alternate model system (*e.g.,* rat) that lacks such a proliferation of paralogs.

Despite insights into EZHIP retention, motifs, expression, and duplication, our analyses did not identify the evolutionary origin of EZHIP. We used both sequence-based (48–50) and structure-based (51) tools to search for distantly related homologs in mammalian and bird genomes. These efforts yielded no convincing matches to any of the conserved EZHIP protein motifs. *EZHIP* could have originated from non-coding regions of the genome, but it is currently not possible to confidently detect such an origin. Thus, the evolutionary origins of this potent PRC2 regulator in mammals remain an open question.

What insights can we glean about the function of *EZHIP* from our analyses of its evolutionary innovation? *EZHIP* arose in placental mammals, whereas the PRC2 complex may date back close to the origin of eukaryotes (52–54). The three primary functions of PRC2 in multicellular eukaryotes are developmental transitions (1, 2), transposon defense (3, 4), and genomic imprinting (2). Of these three roles, genome imprinting is the only function that occurs exclusively in placental mammals, although all three processes vary between animal lineages. Imprinting is an epigenetic process in which gene expression depends on whether a gene is inherited from the mother or the father. Pioneering work from Robert Trivers and David Haig has suggested that this process may have evolved in placental mammals due to an epigenetic tug-of-war between the paternal and maternal genomes over resource allocation to the embryo *in utero* (55). Given its recurrent innovation, evolutionary origin in placental mammals, and germline-enriched expression, we hypothesize that *EZHIP* arose and continues to evolve rapidly to regulate genome imprinting in placental mammals. Indeed, two recent studies directly implicate EZHIP in imprinting function in mice (38, 56). Factors functioning in interparental parental conflict over resource allocation, rather than more critical germline functions, would be predicted to tune but not abolish maternal imprints (such as H3K27me3). This may explain why *EZHIP* deletion mutants do not show severe fertility defects.

The evolutionary characteristics of *EZHIP* – placental mammal-specific origin, X-chromosomal localization, germline-specific expression, rapid sequence divergence, and dynamic gene duplication – are highly reminiscent of the short histone H2A histone variants that arose and proliferated on the X chromosome of placental mammals (57). We previously showed that one short H2A histone variant, H2A.B, functions as a biparental-effect gene, with paternal and maternal contributions synergistically promoting embryo viability and *in utero* growth (58). Like short histone H2A variants, particularly H2A.B, *EZHIP* exhibits evolutionary signatures characteristic of genes mediating genetic conflict between maternal and paternal genomes. Both short H2A variants and *EZHIP* originated exclusively in placental mammals, coincident with the evolution of invasive placentation. Both are X-chromosomal, which could facilitate the rapid evolution of parent-of-origin effects that mediate conflicts over resource allocation to offspring through hemizygous selection in males (59–62). The recurrent duplication and divergence of *EZHIP* across mammalian lineages (including 11 X-chromosomal *Ezhip* paralogs in mice) mirrors the extraordinary evolutionary dynamism of short H2A variants, including up to 20 H2A.L genes in mice (58). Finally, *EZHIP’s* accelerated evolution parallels that of the H2A.B and H2A.P variants, suggesting that *EZHIP* participates in a molecular arms race, as expected for genes mediating genetic conflict, where paternal and maternal contributions have different optimal states.

The multiple parallels between short histone H2A variants, such as H2A.B, and *EZHIP* further suggest that *EZHIP’s* primary function may be to mediate interparental conflict over embryonic development *in utero* in placental mammals. *EZHIP’s* germline-restricted expression and its role in regulating H3K27me3, which can mark imprinted genes (63) that control growth and resource allocation, could place it directly at the interface between parental genomes. Our findings that the primate-specific *EZHIP2* paralog is expressed exclusively in the male germline, whereas *EZHIP* is expressed in both ovary and testis, suggest these paralogs may have functionally diverged and now carry out distinct roles in mediating paternal versus maternal influences on offspring development, with paternally expressed genes predicted to favor increased maternal investment and maternally expressed genes predicted to balance investment across all offspring. Since imprinted loci vary among mammalian species and even among closely related murid and primate species (39–47), it is tempting to speculate that the evolutionary innovation of *EZHIP* may facilitate differential regulation of imprinted loci in placental mammals.

A role for *EZHIP* in mediating interparental conflict over resource allocation to offspring may also partially explain its rare, idiosyncratic losses in multiple Afrotherian species. Elephants possess components of the PRC2 complex and have detectable H3K27 methylation (64, 65). Therefore, the loss we observe in *Afrotheria* is unlikely to reflect a global loss of PRC2 function or components. In these species, inter-parental conflict over fetal resource allocation may either be resolved, mitigated, or accommodated by alternative mechanisms, such as changes in placental structure, imprinting architecture, or other PRC2 regulators, in the absence of *EZHIP*. Indeed, *EZHIP* might be only one of several possible molecular solutions to parental conflict. Its retention might be favored in lineages where PRC2 inhibition is beneficial, but dispensable in lineages where life-history traits, reproductive strategies, or chromosomal context (e.g., differences in X-linked gene content and dosage compensation) make alternative solutions more favorable. Together, *EZHIP* losses in Afrotheria, its rapid evolution under diversifying selection in other mammalian lineages, and its recurrent duplication across lineages highlight that similar genetic conflicts often yield different modes of innovation rather than a single obligate outcome.

## Methods

### Identification of *EZHIP* orthologs

To identify *EZHIP* orthologs, we iteratively queried the genomes of more than 80 placental mammals (see species list in Table S1), representing primates, Glires, ungulates, carnivores, bats, Afrotheria, and Xenarthra. We also queried two non-placental mammalian outgroup species: the gray short-tailed opossum (*Monodelphis domestica*) and the platypus (*Ornithorhynchus anatinus*). We used TBLASTN (49, 66) to perform a homology-based search, using human EZHIP as the initial query. We used genomes from NCBI’s non-redundant nucleotide collection (nr/nt) and whole-genome shotgun contig (wgs) databases to query for homology.

In some species, especially Glires, a homology search using human EZHIP yielded no results due to poor homology. In these cases, we identified *EZHIP* orthologs using two complementary approaches. First, we conducted syntenic analyses to identify the genomic neighborhood in which *EZHIP* is expected to be located. Second, we used EZHIP sequences from more closely related species as TBLASTN queries to identify *EZHIP* orthologs; for mouse, we used existing literature (12). Using the identified orthologs, we re-queried the genome to detect any paralogs. For example, using mouse EZHIP to query the mouse genome led us to identify 15 additional homologous EZHIP sequences. In all cases, paralogs were reciprocally blasted against *Homo sapiens* or other closely related species to determine if the identified paralogs were indeed EZHIP-related. Furthermore, we included all orthologs and paralogs with intact open reading frames in our maximum-likelihood phylogenetic analyses described below to confirm their relationship to EZHIP.

We examined the shared synteny (conserved genetic neighborhood) of EZHIP to confirm orthology and to date its origin. Syntenic analyses were performed in two non-placental mammalian outgroup species (*Monodelphis domestica* and *Ornithorhynchus anatinus*) and 15 placental mammals: human (*Homo sapiens)*, mouse (*Mus musculus)*, rat (*Rattus norvegicus)*, guinea pig (*Cavia porcellus*), rabbit (*Oryctolagus cuniculus)*, pig (*Sus scrofa*), sheep (*Ovis aries)*, cow (*Bos taurus)*, horse *(Equus caballus)*, south-central black rhinoceros (*Diceros bicornis)*, cat *(Felis catus)*, dog (*Canis lupus familiaris)*, greater horseshoe bat (*Rhinolophus ferrumequinum*), Asian elephant (*Elephas maximus)*, and armadillo (*Dasypus novemcinctus*). Using the UCSC Genome Browser (67), genomic neighborhoods were identified for all our hits. *EZHIP* is present on the X chromosome, which is more susceptible to evolutionary turnover. Therefore, we identified well-conserved flanking genes surrounding *EZHIP* (*BMP15, NUDT10/11, MAGED1*) using the Genomicus browser (68). Using TBLASTN, the flanking genes were used as queries to identify syntenic regions across the 17 mammalian genomes (Fig. 1B). In some cases, neighboring genes were found many kilobases away from *EZHIP*. However, given the rapidly evolving nature of the X chromosome, we still counted these as syntenic regions if they were found on the same chromosome or scaffold despite their large genomic distance from *EZHIP*. In cases where the syntenic location was split across multiple scaffolds, we indicate this break by double slashes in Fig. 1.

*EZHIP* orthologs were identified based on sequence homology to known EZHIP sequences (predominantly human or rodent), conserved synteny, and phylogenetic relationships (below). Sequences that did not meet these criteria were classified as paralogs or pseudogenes. Pseudogenes were annotated based on the presence of multiple stop codons, frameshifts that disrupt open reading frames, or the absence of any initiating methionine (Table S1).

### Phylogenetic analyses

We performed phylogenetic analyses to resolve relationships among all identified EZHIP homologous sequences and to distinguish orthologs from paralogs. We used the MUSCLE algorithm (Edgar, 2004) in Geneious Prime 2025.0.3 (https://www.geneious.com) to perform protein alignments. Positions present in fewer than half of all sequences were removed from the alignment (de-gapped) to allow for a more stringent analysis, and pseudogenes were also excluded (Data S2 and S7). Phylogenetic trees were constructed using maximum-likelihood methods implemented in PhyML (20, 21) and the JTT substitution model (69) with 100 bootstrap replicates (Fig. 1C, Fig. S4, Fig. S13). PHYML treats the remaining alignment gaps as unknown characters.

### Calculating the rate of protein divergence

We calculated pairwise protein sequence identities for EZHIP across representative species of primates, carnivores, ungulates, and rodents, relative to human, dog, goat, and mouse orthologs, respectively. We similarly calculated pairwise protein sequence identities for primate and rodent EZH2 orthologs (Fig. 3; Table S3). Median species divergence times were obtained from the TimeTree database (www.timetree.org (29)).

### Analyses for evolutionary selective pressures

We analyzed selective pressures on EZHIP and EZHIP2 across diverse mammals and simian primates using the codeml algorithm from the PAML suite (31) (Data S4, S5, and S9). We generated and manually refined codon-based alignments using either the translation align tool in Geneious Prime 2025.0.3 (https://www.geneious.com), or the MACSE algorithm (70). We manually trimmed alignments to remove gaps or regions unique to a single species, as well as poorly aligned regions in the repetitive region of EZHIP. The resulting alignment was used to construct a phylogenetic tree using PhyML with maximum-likelihood methods and the GTR substitution model (20).

We assessed gene-wide purifying selection (Table S2) using codeml model 0. This model assumes a single evolutionary rate across all lineages present in the sequence alignment. We compared the likelihoods between model 0 with dN/dS fixed at 1 (neutral evolution) and model 0 with dN/dS estimated from the alignment. Statistical significance was determined by comparing twice the difference in log-likelihoods between the two models to a χ² distribution with one degree of freedom (31).

To test for site-specific positive selection (Table S4), we compared two pairs of PAML NSsites models. We compared log likelihoods between model 8 (which encompasses 10 codon categories with dN/dS values between 0 and 1 and an additional category with dN/dS > 1) and either model 7 (which has only the 10 categories with dN/dS between 0 and 1) or model 8a (which fixes the additional category at dN/dS = 1). Statistical significance was determined by comparing twice the difference in log-likelihoods between the models to a χ² distribution, with degrees of freedom corresponding to the difference in the number of model parameters (31). We categorize positively selected sites as those that have a Bayes Empirical Bayes (BEB) posterior probability greater than 90% in model 8. We also ran each PAML analysis with four sets of alternative starting parameters (codon models 2 or 3; starting omega 0.4 or 3); our results were robust to these alternative parameters. To validate our findings, we conducted independent analyses using the FUBAR algorithm (33) in Datamonkey (https://www.datamonkey.org), which estimates selection at individual sites and across the entire gene (Table S4).

Gene conversion/recombination can sometimes result in false positive evidence for positive selection (71). To test for this, we ran the GARD algorithm (HyPhy package) (72) on the primate EZHIP and EZHIP2 alignments, specifying 3 rate categories. GARD found some evidence that recombination might have occurred in EZHIP (evidence ratio 85.4, with one breakpoint) but not EZHIP2 (evidence ratio 0). We therefore divided the EZHIP alignment at the predicted recombination breakpoint and reran PAML on both segments. Positive selection is still reported on both segments. Because the GARD evidence ratio is somewhat weak (evidence ratios of <100 may reflect rate variation within the alignment instead of true recombination), we choose to report positive selection results only for the original, full-length alignment.

### Motif analyses

Ten motifs were identified using both MEME and MAST (23) on a single representative copy of EZHIP from more than 70 placental mammals (Data S3). While motifs 1-4 and 8-10 were typically found only once in each EZHIP homolog, motifs 5-7 were often repeated several times in each protein. In addition, they showed high sequence similarity to one another, as reported by MAST analyses. Therefore, we excluded motifs 5-7 from the initial analyses to avoid erroneous conclusions about retention or loss and instead analyzed them for tandem repeats (see below). All identified motifs had E-values < 10^-5^. We used custom R code, utilizing the universal motif (73) and ggseqlogo (74) packages, to identify residues within each motif that are conserved in at least 80% of motif instances, and to replot motifs with our chosen color scheme. In cases where neither program detected motifs, all sequences within a given clade (e.g., carnivores) were aligned to manually confirm the absence of a motif. If the pairwise identity exceeded 50%, we manually classified this motif as present. In addition, the entire human EZHIP sequence was used to identify a nuclear localization sequence (NLS) using PSORT (26) (https://psort.hgc.jp/). This NLS overlaps with a well-conserved stretch in motif 5.

We used the identified motifs to perform more sensitive homology searches of genomes and proteomes from selected species. We used three methods for these searches: the FIMO and MAST algorithms (MEME suite), and Hidden Markov Model (HMM) searches. Code and output is provided at https://github.com/jayoung/EZHIP_evolution/tree/main/motif_analysis/protein_motif_databas e_searches_process_output.md.

The species and genome assemblies searched were human (hg38); four Afrotheria lacking KLP, comprising African and Asian elephants (loxAfr3 and mEleMax1), dugong (mDugDug1) and manatee (TriManLat1.0); three marsupials, comprising wallaby (mMacEug1), opossum (mMonDom1) and Tasmanian Devil (mSarHar1.11); and platypus (mOrnAna1). The databases for each search were (a) annotated protein sequences in each genome, obtained via the NCBI ftp site, with any selenocysteine residues (U) converted to cysteine residues because they are not tolerated by the MAST algorithm (b) 6-frame translations of each complete genome, generated using the EMBOSS package getorf (parameters -minsize 300 -find 0), and cleaned so that stretches of >12 X characters - the result of translating genome assembly gaps - were replaced by 9 Xs (long X stretches caused false positive matches in MAST searches).

MAST searches were performed with default parameters, and output was parsed in R. For the HMM searches, we first used R code to parse MEME output and then exported fasta files of all instances of each motif among the original input EZHIP sequences. We then used hmmbuild and hmmpress (HMMER package, version 3.4) to convert fasta files to HMMs. We then used each motif HMM to search the 6-frame genome translations and NCBI proteome databases listed above, with the hmmsearch algorithm (--max parameter for extra sensitivity). Output was parsed in R.

We analyzed *EZHIP* sequences from 50 species using Tandem Repeat Finder (TRF) (https://tandem.bu.edu/trf/trf.html) (27) to assess repeat length, copy number, and sequence similarity. For each sequence, non-overlapping repeats were annotated based on the minimum repeat length. We further annotated repeat sequences to ensure they overlapped with the open reading frame. We then used iterative alignments between humans and each species, or between species within each clade, to annotate and classify repeats as mammal-wide (suggesting an origin in the last common ancestor of all mammals), clade-specific, or species-specific. For *EZHIP2*, TRF could only identify repeats if less stringent parameters were used (alignment parameters = “2,3,5”, minimum score = 30). Motif logo plots (Fig. 2C, D, and Fig. S9) were created from these alignments using WebLogo (<u>weblogo.berkeley.edu</u>) (75).

### RNA-seq analyses

We analyzed publicly available transcriptome data from human tissues (76, 77), rhesus macaque tissues (76), human spermatogenesis (78), and human embryos (79) to quantify the expression of *EZHIP* and *EZHIP2* (87.4% identical to EZHIP) (Table S5). We downloaded FASTQ files using NCBI’s SRA toolkit (https://www.ncbi.nlm.nih.gov/books/NBK158900) and mapped reads to species-specific genome assemblies using STAR (80) with the options “–outMultimapperOrder Random – outSAMmultNmax 1 –twopassMode Basic” to randomly assign multimapping reads to a single location. Gene-level read counts were obtained using genomic coordinates of each ORF and BEDTools multicov (81). RPKM values were calculated by dividing read counts by the total number of mapped reads (in millions) and transcript length (in kilobases).

To evaluate *EZHIP* and *EZHIP2* expression in cancers, we obtained RNA-seq BAM files (n ≈ 11,000) from The Cancer Genome Atlas (TCGA) Genomic Data Commons (GDC) as aligned, coordinate-sorted “rna_seq.genomic.gdc_realn.bam” files together with the corresponding metadata. To quantify reads overlapping a single custom locus (e.g., *EZHIP2*), we constructed a minimal gene annotation in GTF format containing a single gene with one transcript and one exon, ensuring that standard GTF syntax, consistent gene_id and transcript_id attributes, and chromosome names matched to the TCGA BAM headers. Read counting was performed using featureCounts from the Subread package (version 2.0.3-GCC-11.2.0) in paired-end mode, counting fragments that overlap annotated exons. For each BAM, featureCounts was run independently with options “-t exon -g gene_id -p -B -C” and an unstranded setting (-s 0), and the custom GTF as the annotation input. Per-sample *EZHIP2* counts were then collated into a single matrix for downstream analyses.

### Transgenic cell line generation

We adapted HEK293T landing pad attP* cells (82) and seeded them to 60% confluency in 6-well plates. The next day, 1.6 μg of attB*-containing *EZHIP* (or *EZHIP2*) plasmid and 0.4 μg of Bxb1 expression vector (pHPHS115) were transfected per well using Lipofectamine 3000 (Thermo L3000008). Cells in each well were expanded into a 10-cm dish, 48 hours post-transfection. Cells were then selected with 2 μg/ml puromycin, added 72 hours post-transfection. Puromycin selection was terminated after 4 days, and successful integration and selection were confirmed by flow cytometry analysis of mNeonGreen-positive cells.

### Western blot analyses

Transgenic cells were seeded in 6-well plates. 24 hours later, we added 2 μg/ml doxycycline to induce *EZHIP* (or *EZHIP2*) expression. Two days post-induction, cells at 70% confluency were harvested with Buffer A and benzonase and immediately stored at -80°C. Frozen whole-cell extracts were thawed by shaking at room temperature. Samples were run on a 12% precast polyacrylamide gel (Bio-Rad) followed by transfer onto a 0.2mm nitrocellulose membrane (Bio-Rad). Membranes were washed with TBS and blocked with Licor TBS blocking buffer. Primary antibodies targeting tubulin (abcam ab6046), H3 (abcam ab1791), or H3K27me3 (Cell Signaling, C36B11) were used, followed by goat anti-rabbit IRDye 680 secondary antibody (Licor 926-68071).

### Data and code access

Supplementary Data files include all sequences used in trees and all mammalian orthologs and paralogs of EZHIP described in the paper. Code used for analysis can be found in a GitHub repository (https://github.com/jayoung/EZHIP_evolution).

## Acknowledgments

We thank Mary C. Rominger for her initial work in identifying some of the primate orthologs and paralogs found in Fig. S11. We thank Dr. Peter Lewis for discussions, and Abby Clyde, Peter Dietzen, Dr. Jeremy Hollis, Dr. Sarah Leichter, and Dr. Ben Martin for comments on the manuscript. This work was supported by grants from the National Institute of General Medical Sciences at the National Institutes of Health: K99/R00 GM149928 (to P.R.), R35 GM139429 (to T.T.), R01 CA262556 (to A.B.), and R01 GM074108 (to H.S.M.), and from the Howard Hughes Medical Institute (to M.T.M. and H.S.M.). The funders played no role in study design, data collection, interpretation, or the decision to publish this study. T.T. holds the David and Deborah Lycette Endowed Chair for Cancer Research and H.S.M. is an Investigator of the Howard Hughes Medical Institute.

---

**Supplementary Figure S1.**
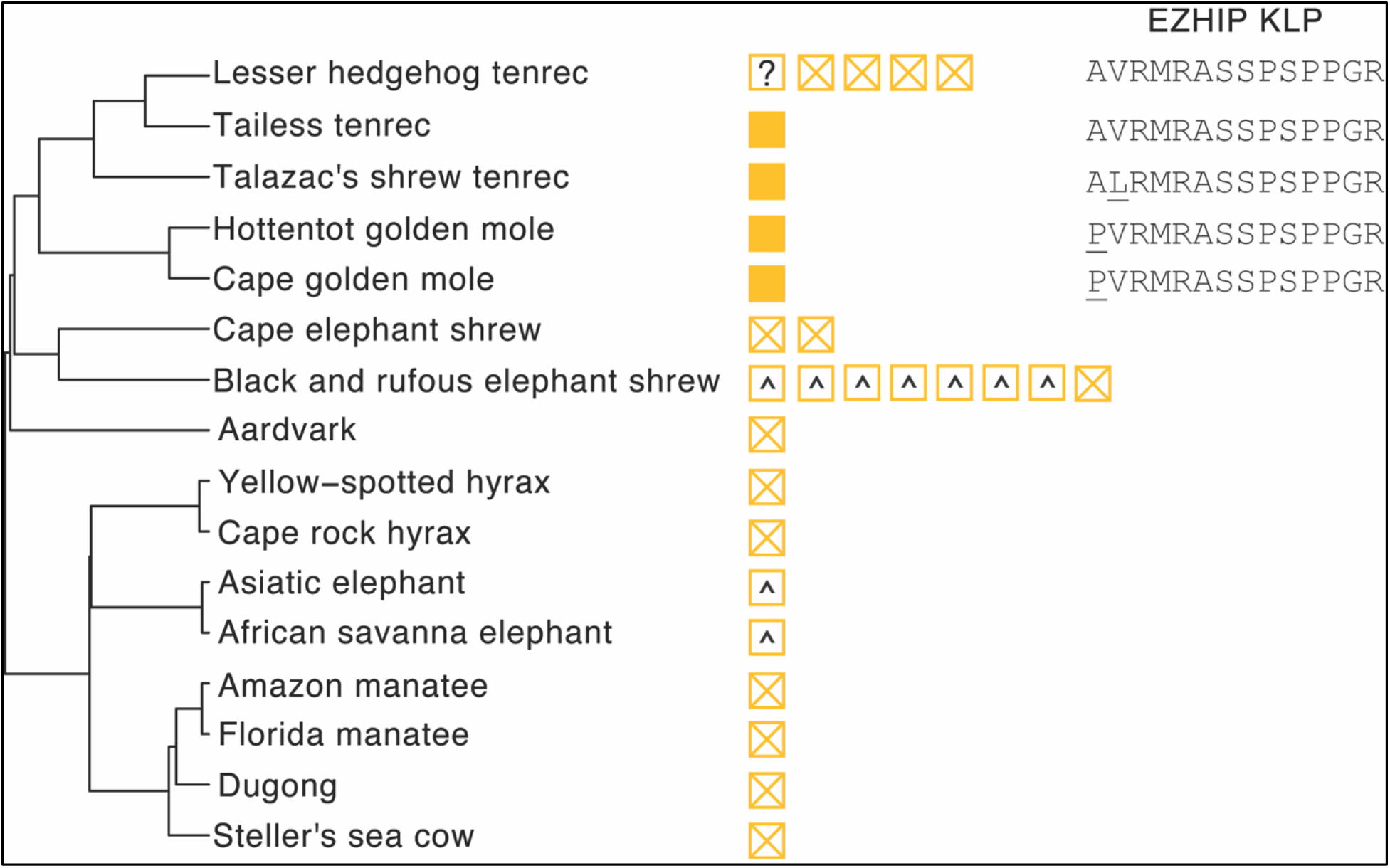
*EZHIP* retention in Afrotherian species. *EZHIP* gene copies (yellow boxes) are illustrated alongside an Afrotherian species tree (22). Filled boxes represent largely intact *EZHIP* sequences. Empty boxes containing an X represent putative pseudogenes, *^* represents a lack of a KLP sequence, andrepresents a frameshift in the open reading frame. The KLP sequence from five *EZHIP* copies is shown, with changes relative to human *EZHIP* underlined.

**Supplementary Figure S2.**
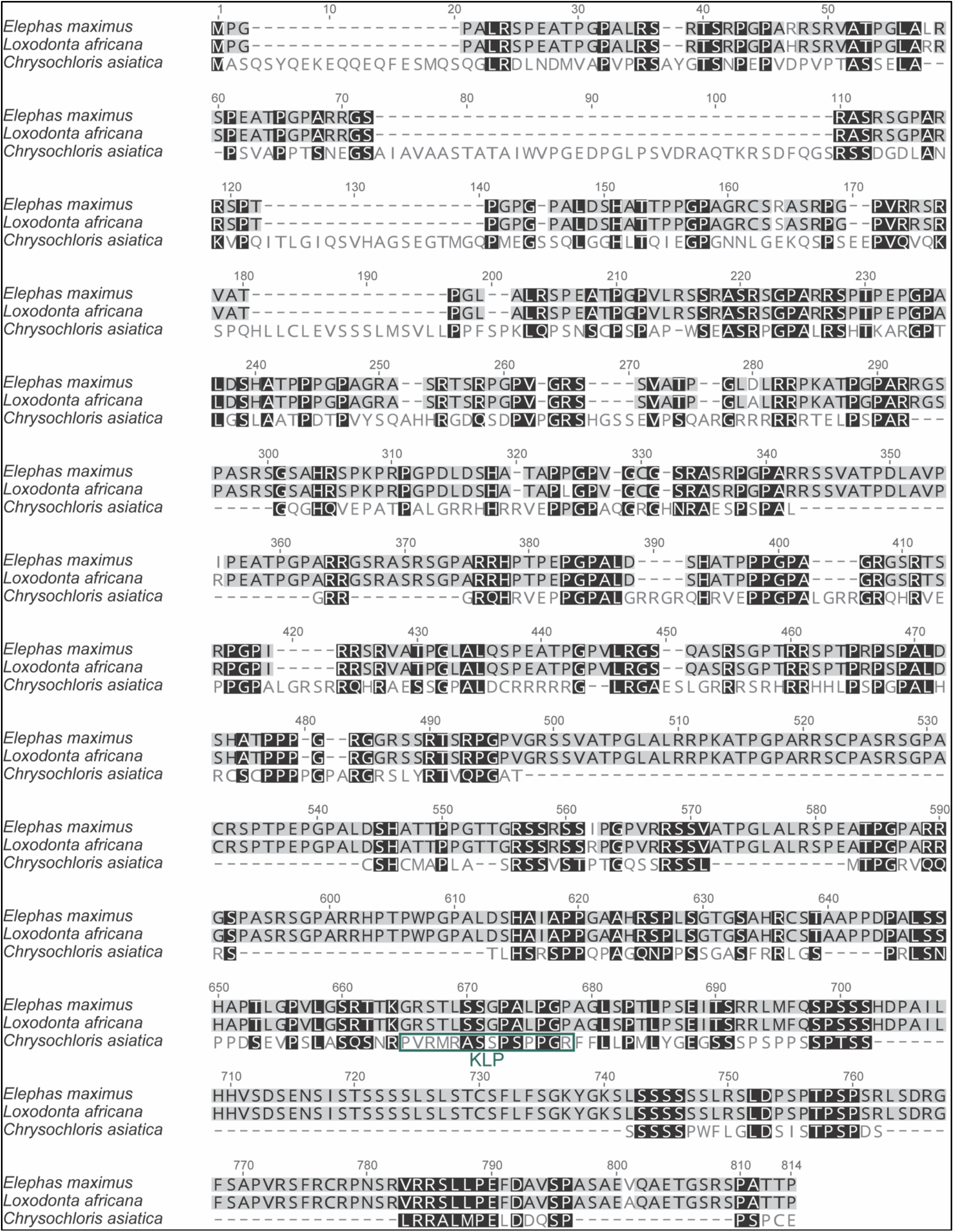
Protein sequence alignment of selected Afrotherian species. EZHIP alignment from two elephant species, *Loxodonta africana* and *Elephas maximus*, and from the cape golden mole, *Chrysochloris asiatica*, is shown. Residues with black and grey backgrounds indicate high sequence similarity and hyphens indicate gaps in the alignment. The KLP motif in the Cape golden mole is indicated with a dark green-outlined box.

**Supplementary Figure S3.**
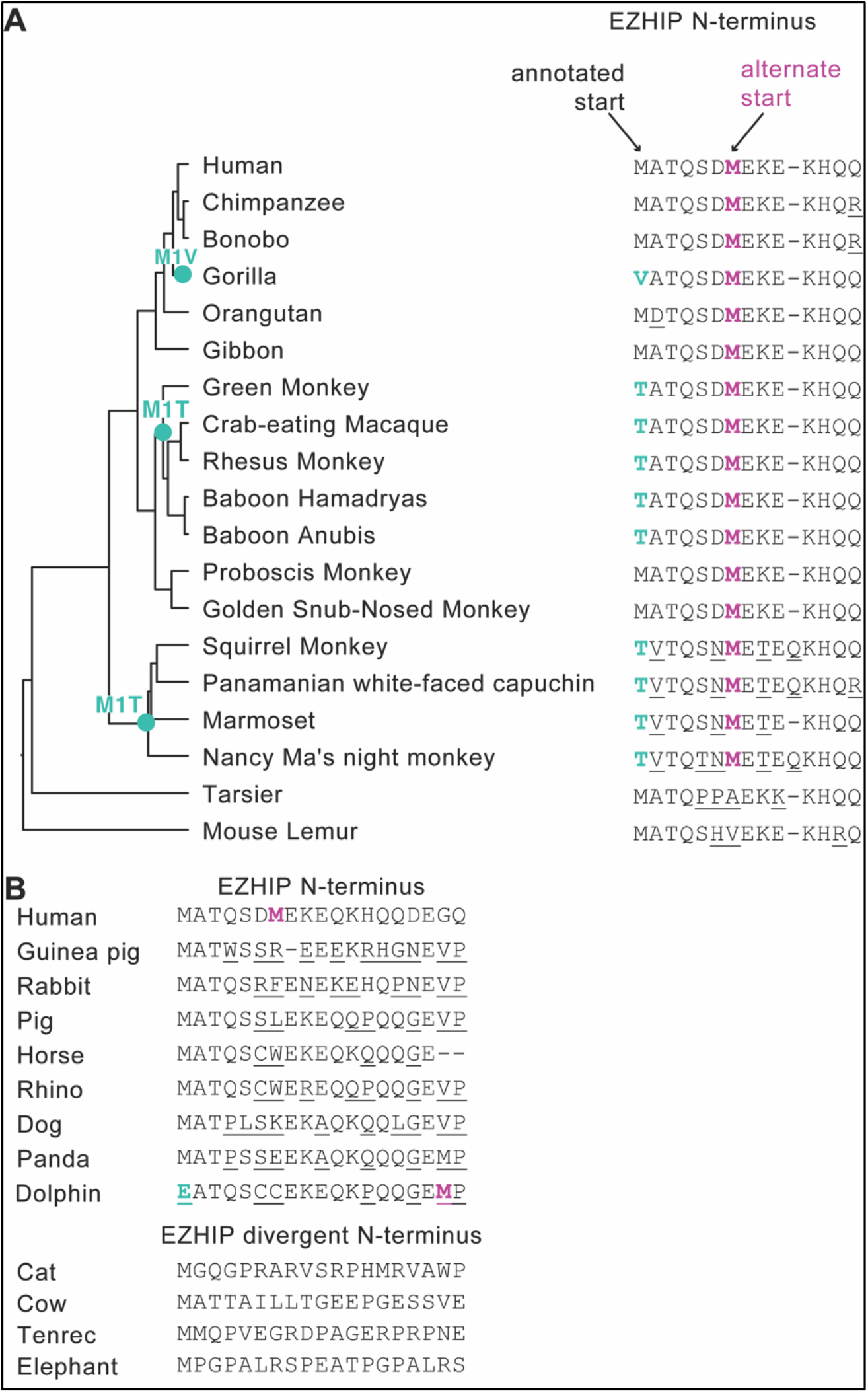
Multiple primates and several additional mammals may have evolved an alternate start codon. **A.** The first 14 amino acids of primate EZHIP are shown beside a primate species tree. Repeated loss of the previously annotated start codon is indicated with a teal dot and the mutation. Hyphens indicate gaps in the alignment. We highlight changes to the previously annotated methionine (teal), a conserved, potentially alternate start (pink) and other differences from the human sequence (underlined). **B.** The first 17 amino acids of EZHIP in representative mammals are shown. Any sequence changes relative to the human sequence are underlined. Alternate human and dolphin start codon are indicated in boldface pink. Cat, cow, tenrec, and elephant have a divergent N-terminal tail that cannot be aligned with confidence.

**Supplementary Figure S4.**
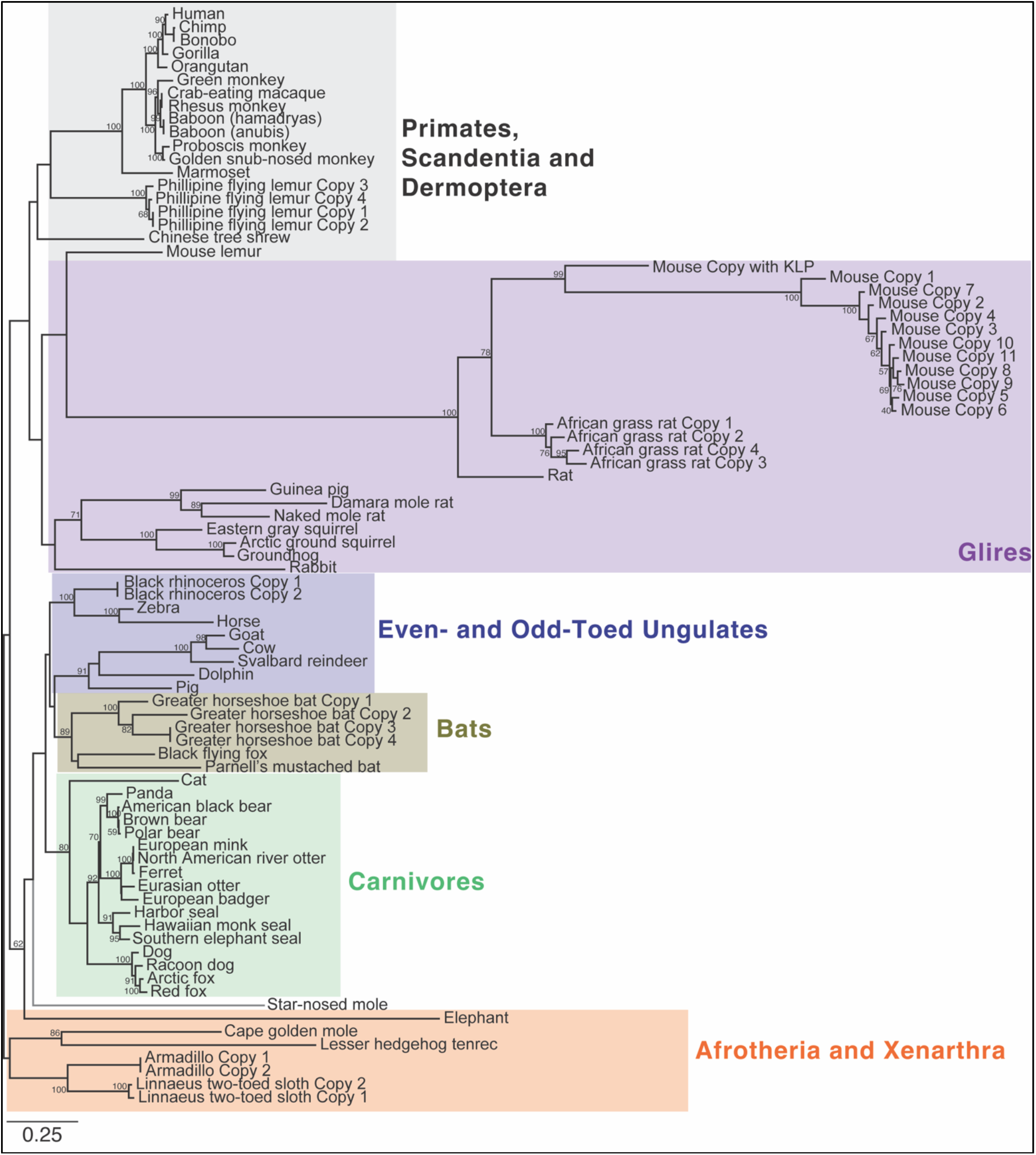
Protein phylogeny of mammalian EZHIP. A maximum-likelihood protein phylogeny of syntenic EZHIP copies with complete ORFs across 64 representative mammalian species (also see Fig. 1C). Colored boxes in the phylogeny indicate different mammalian lineages – primates (gray), *Glires* (purple), carnivores (green), ungulates (blue), bats (brown), and Afrotheria and Xenarthra (orange). Bootstrap values greater than 50 are shown alongside the node they represent. The bottom-left scale bar below the phylogeny represents 0.25 substitutions per site.

**Supplementary Figure S5.**
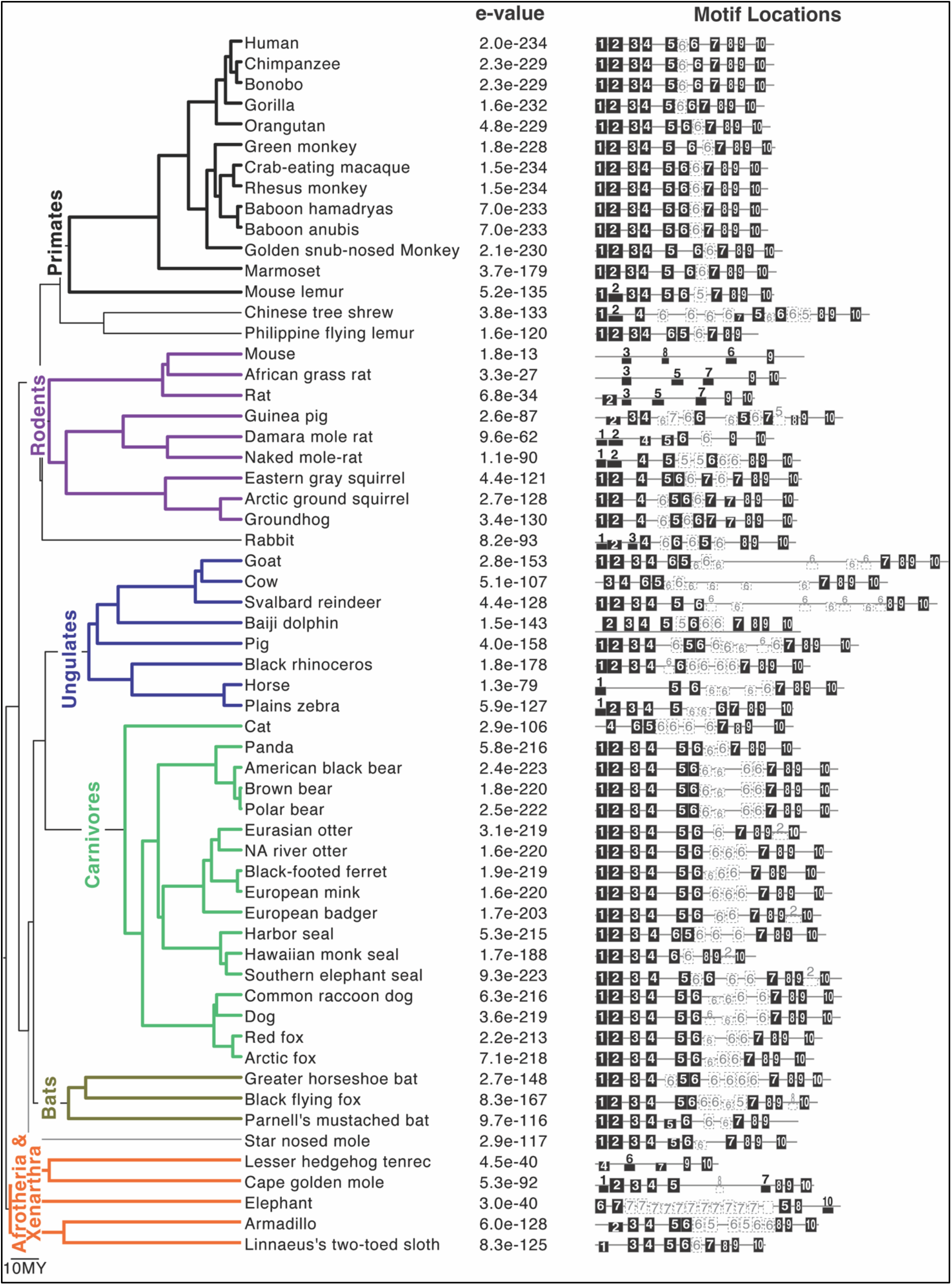
EZHIP has seven conserved motifs and tandem repeats. A mammalian species tree (colored similar to Fig. 1C) is shown alongside a representative protein schematic with identified motifs that are typically found as one copy per gene (e.g., black filled boxes with numbers 1-4 and 8-10) or in multiple copies (e.g., motifs 5-7 in empty dashed boxes). Motif heights represent the significance of a motif site within the sequence, with taller motifs representing more statistically significant sites. E-values from MAST analyses are indicated to the left of motif schematics. See Methods for details.

**Supplementary Figure S6.**
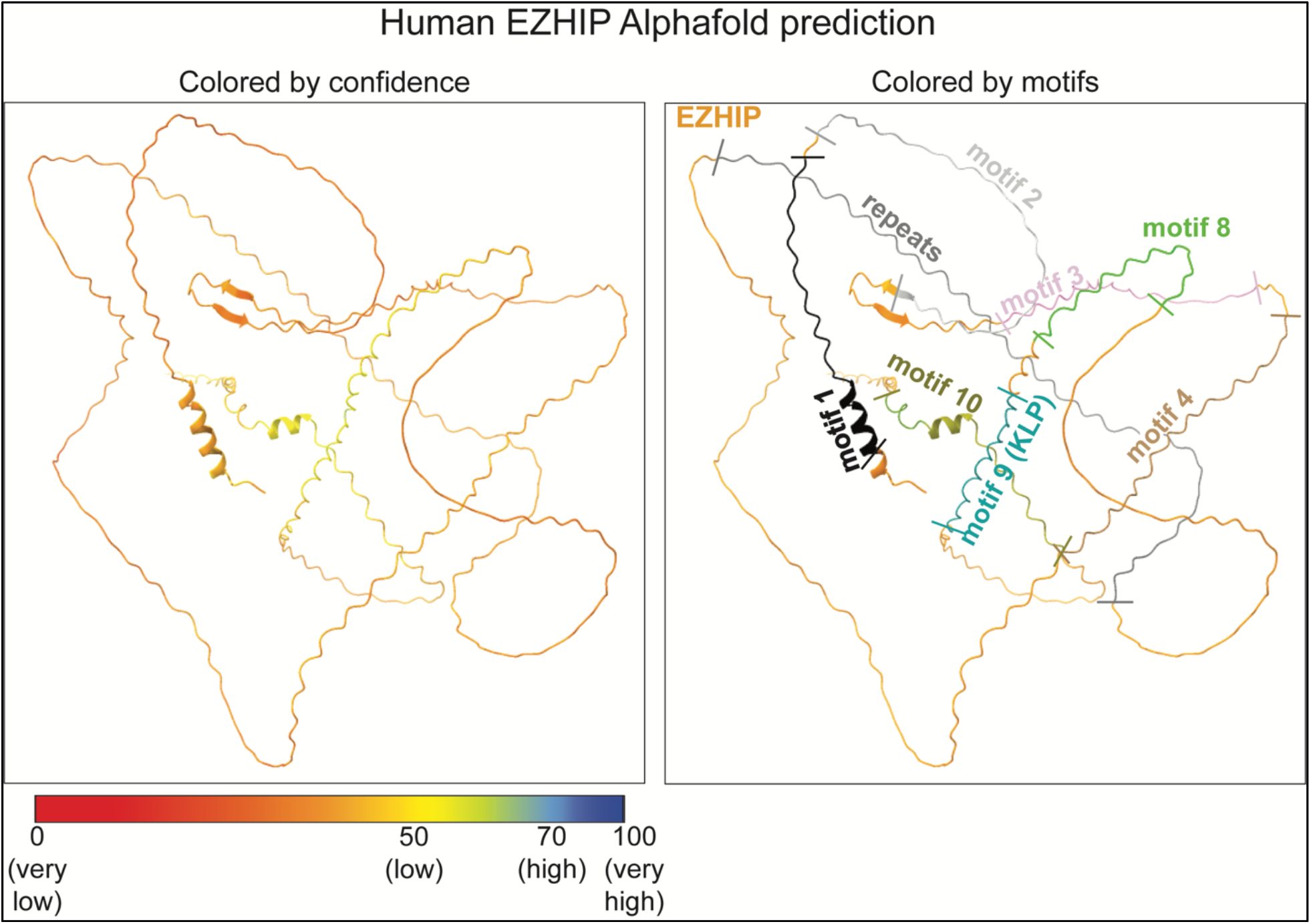
Predicted structure of EZHIP. AlphaFold predictions of EZHIP colored to show confidence in predictions (left), or to highlight motifs identified in our study (right).

**Supplementary Figure S7.**
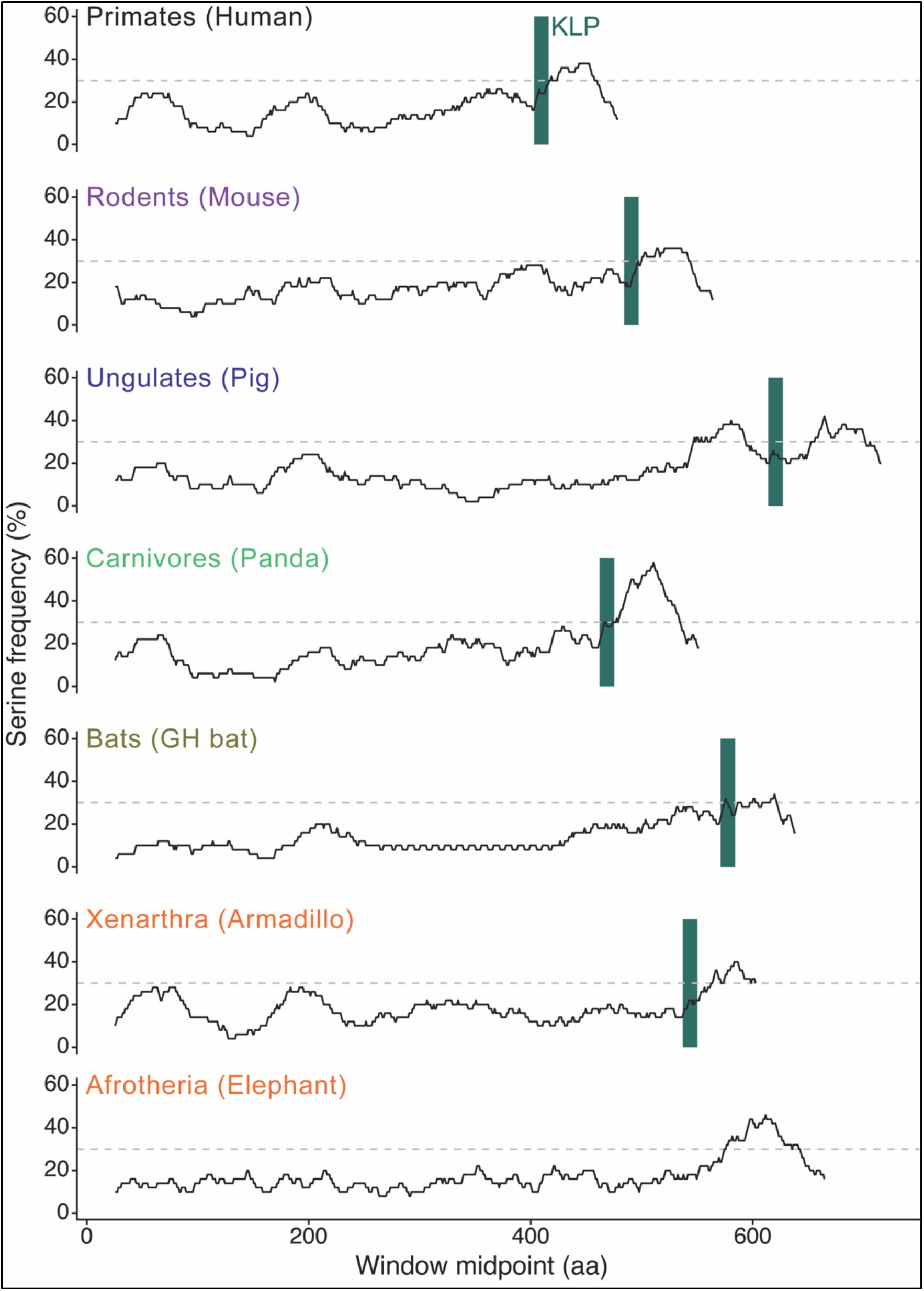
EZHIP has a serine-rich region downstream of the KLP motif. We plot the percentage of serine residues in a 50 amino acid sliding window, sliding in 1aa increments, across EZHIP proteins from representative mammalian species. The KLP is highlighted with a green vertical box.

**Supplementary Figure S8.**
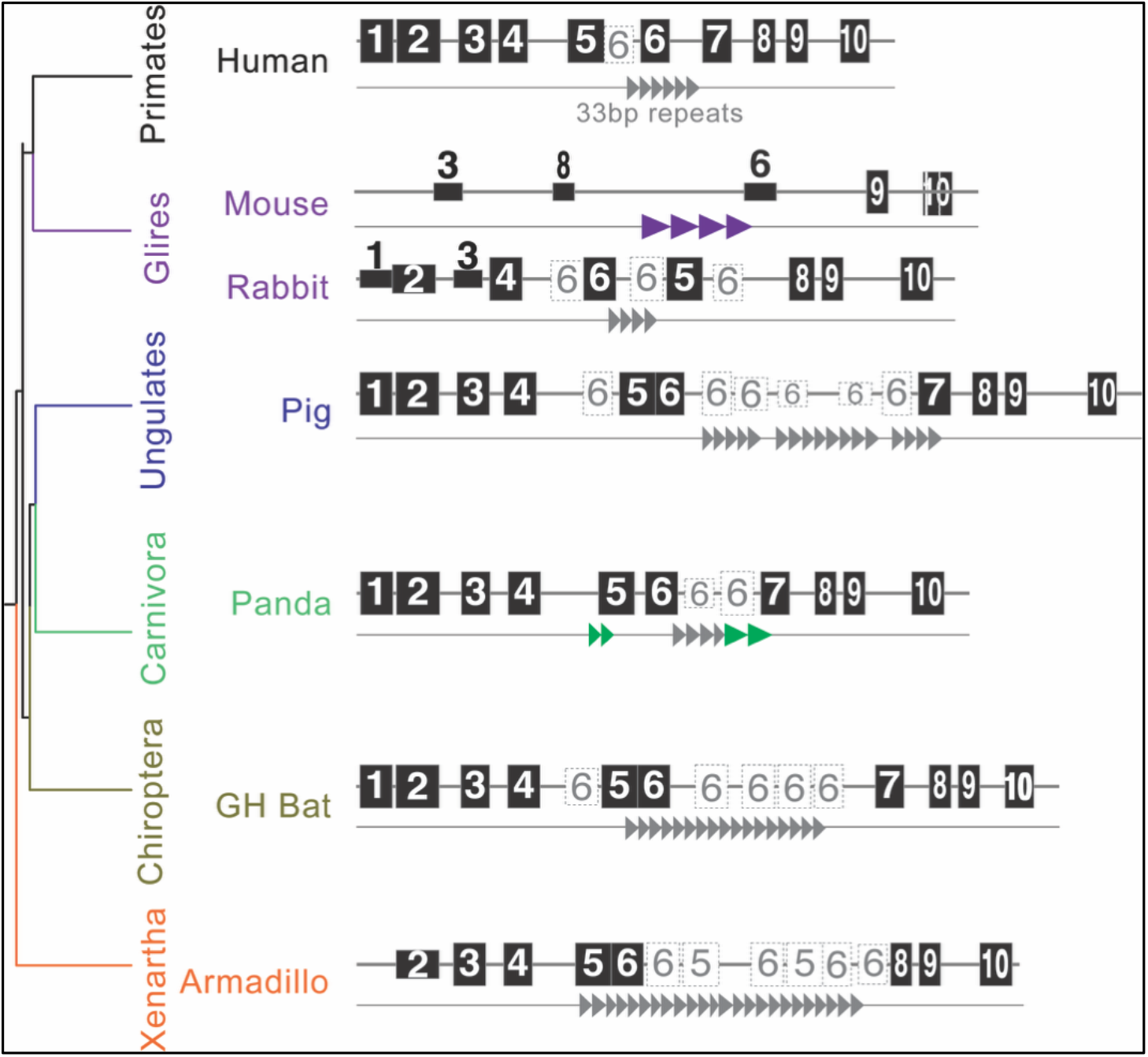
EZHIP motifs 5-7 include tandem repeats. A simplified species tree of representative mammalian species groups alongside selected EZHIP protein schematics (motifs are indicated by black filled boxes with numbers and drawn to scale) represented as in Fig. 2B. Repeats detected in EZHIP using Tandem Repeat Finder (TRF) are represented as triangles and indicate overlap with motifs 5-7 identified by MEME analyses. Most *EZHIP* orthologs had varying numbers of an ancient, common 33-bp tandem repeat (gray triangles), or lineage-specific repeats (purple in mouse and green in panda).

**Supplementary Figure S9.**
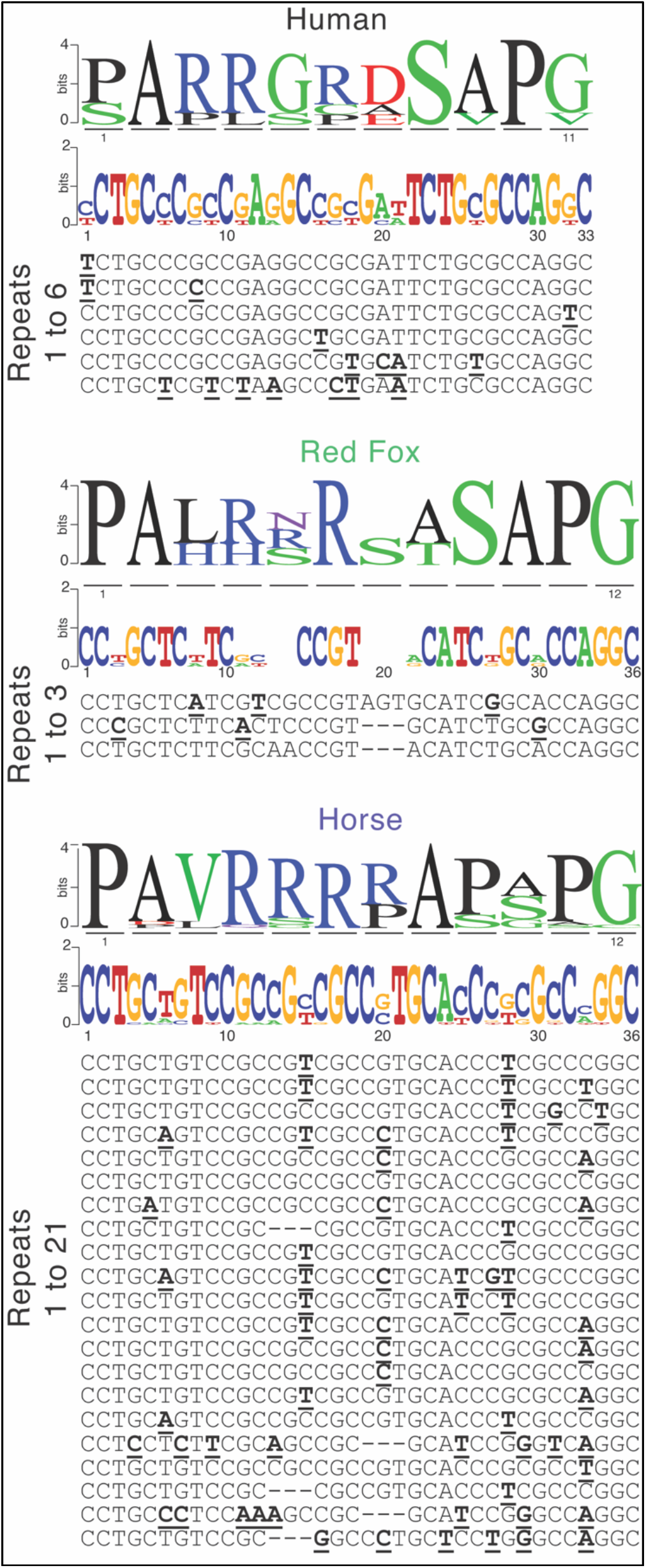
Repeats are conserved within mammalian species. Amino acid and nucleotide logo plots with alignment between copies of tandem repeats found within *EZHIP* of human, red fox, and horse. Hyphens indicate gaps in the alignment. Residues different from the first repeat unit are in boldface and underlined. Logo plots as in Fig. 2.

**Supplementary Figure S10.**
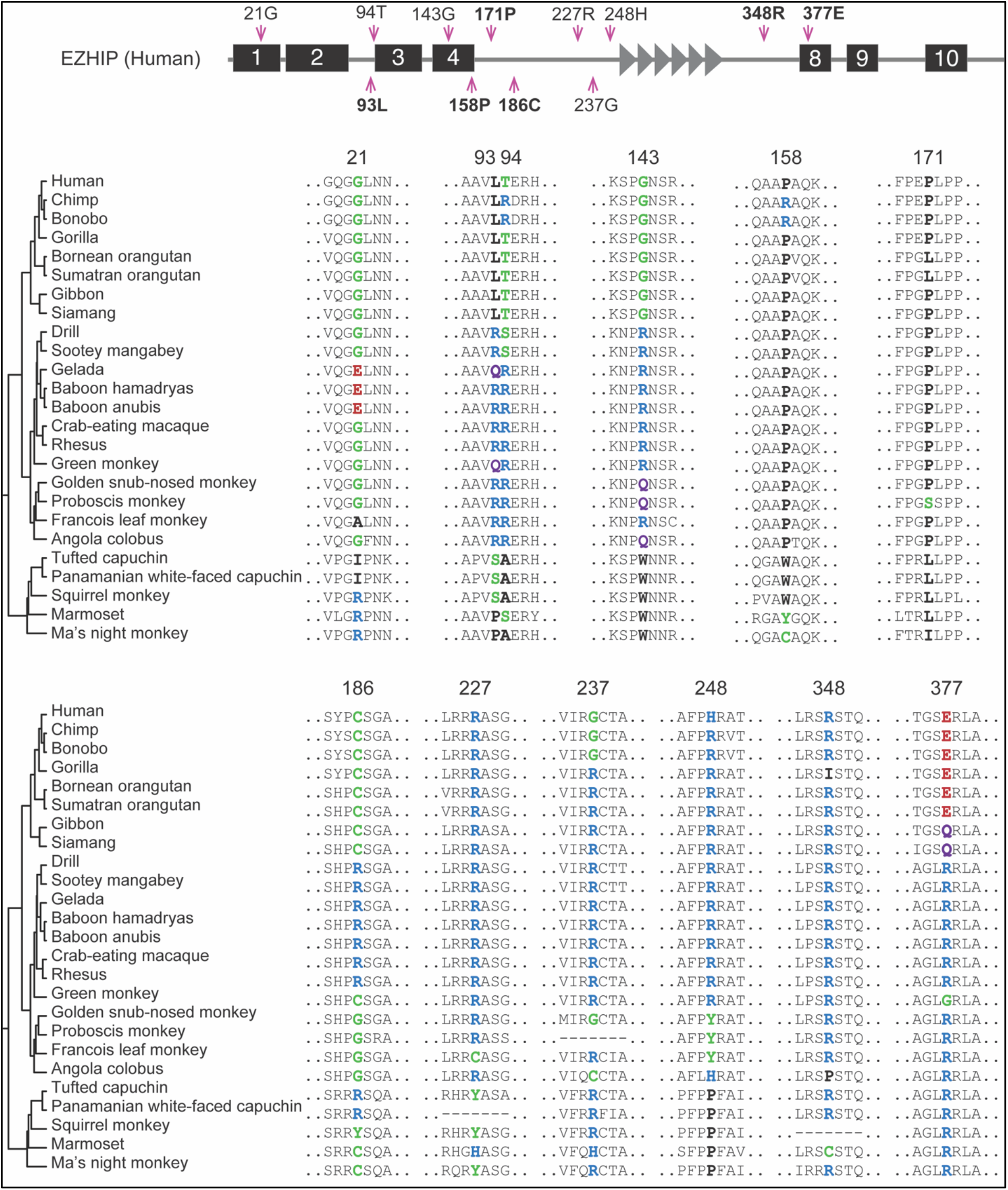
Evidence for positive selection at a subset of sites across simian primate EZHIP. Alignments of regions surrounding positively selected sites (colored, bolded amino acid residues) in EZHIP across simian primates identified by PAML and FUBAR (see Fig. 3B). Hyphens indicate gaps in the alignment. Colors of residues highlight their biochemical properties: hydrophobic (black), negatively charged (blue), positively charged (red), polar (green), and others (purple).

**Supplementary Figure S11.**
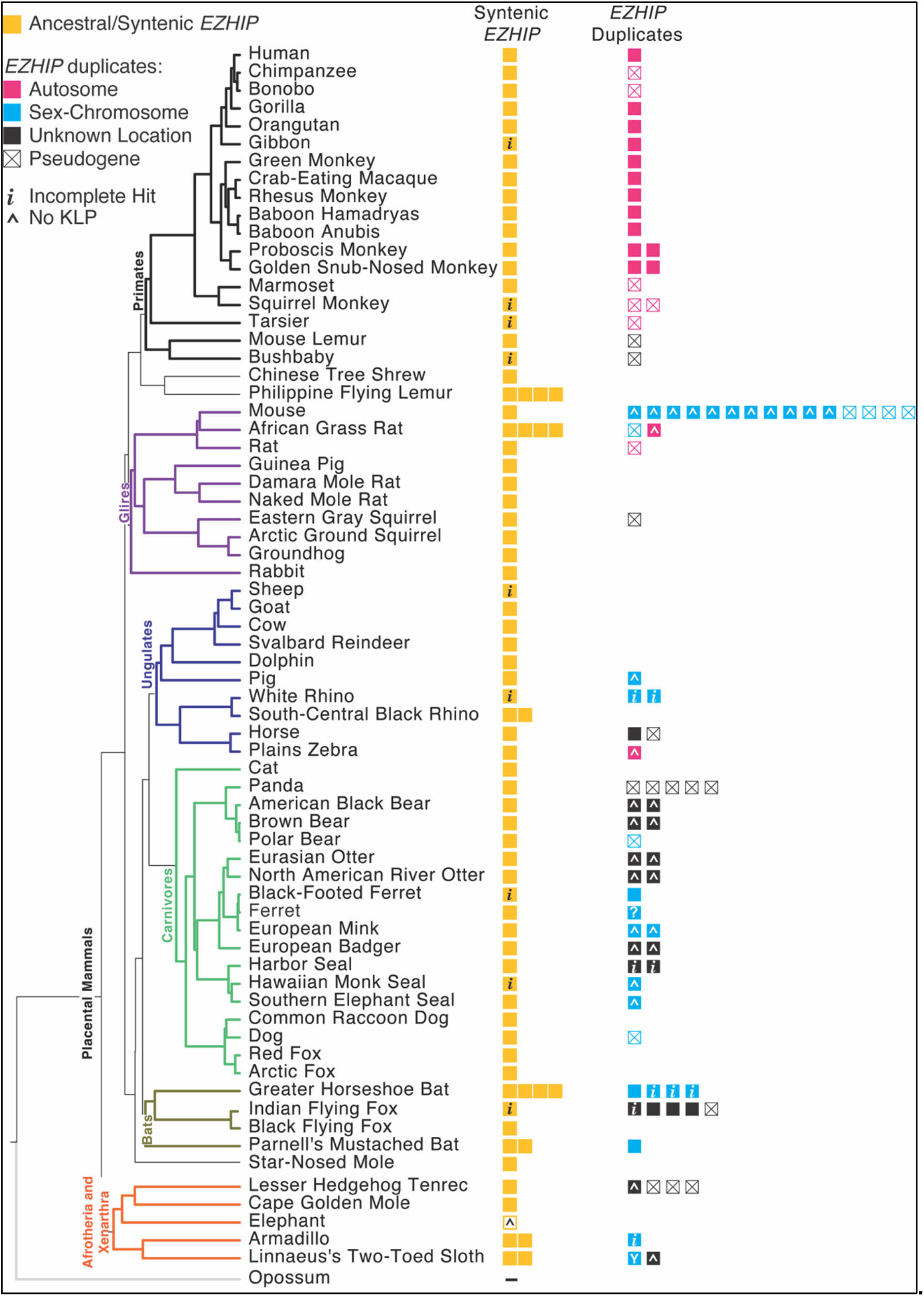
*EZHIP* has repeatedly duplicated in mammals. *EZHIP* copies are represented as colored boxes alongside a mammalian species tree (22) (colored as in Fig. 1C). Orthologs and duplicates in the same syntenic location are shown as yellow-filled boxes. *EZHIP* duplicates can be found on autosomes (pink boxes), sex chromosomes (blue boxes), or at unknown chromosomal locations (black boxes). Most sex chromosome duplicates are found on the X chromosome, except for the duplicate in Linnaeus’s two-toed sloth that is found on the Y chromosome. Boxes containing an X represent putative pseudogenes, *i* represents incomplete sequences due to gaps in genome assembly, and ^ represents a lack of a KLP sequence. In one case, a ferret duplicate, we are unable to confirm if the sequence has coding potential due to variability between genome assemblies; therefore, it is represented with a ?.

**Supplementary Figure S12.**
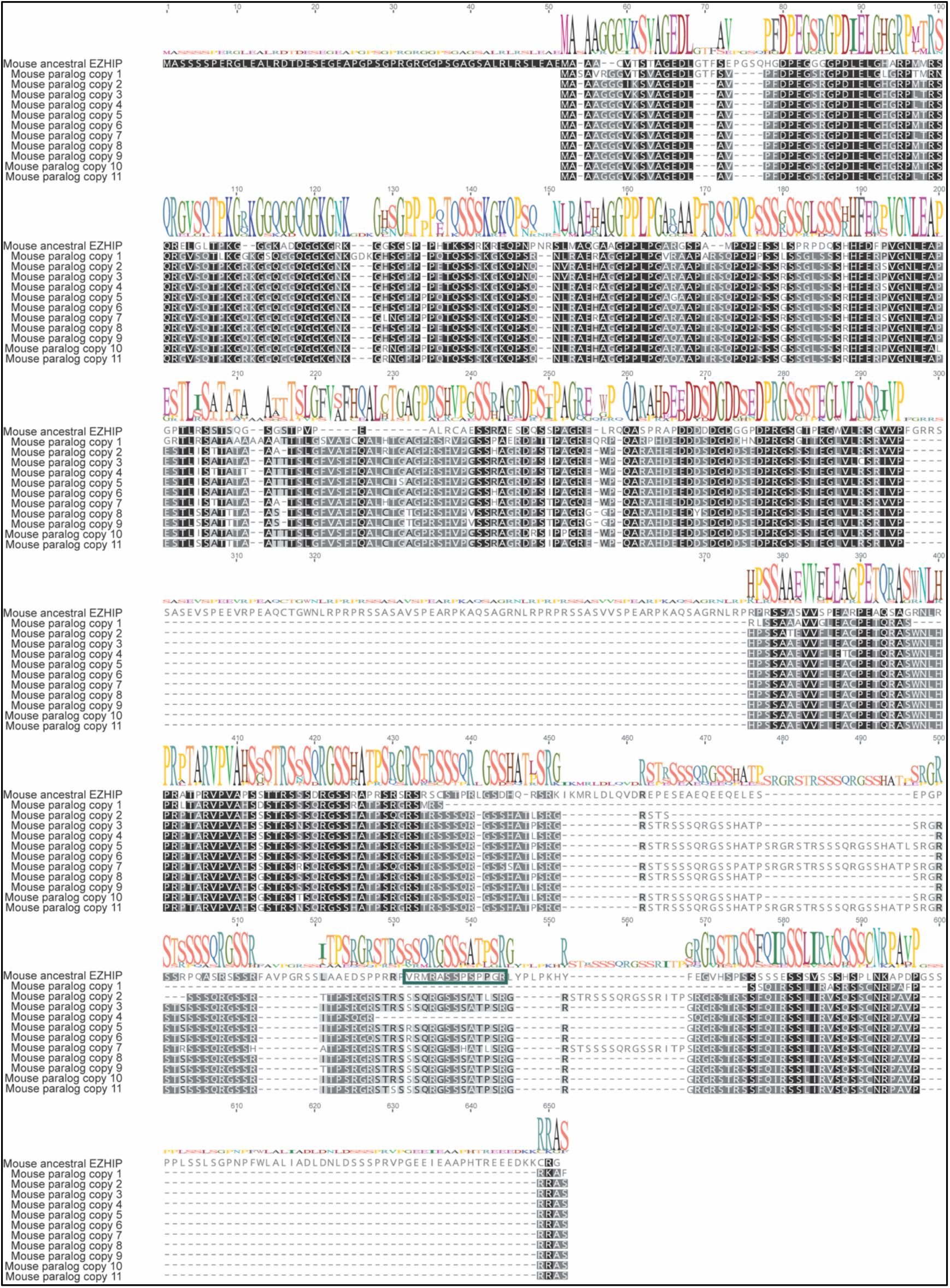
Mouse carries 11 EZHIP paralogs that lack the KLP. The mouse Ezhip ortholog is aligned to 11 Ezhip paralogs identified in the mouse genome. Hyphens indicate gaps in the alignment. Residues with black and grey backgrounds indicate high sequence similarity, and a logo plot showing sequence conservation is displayed above the alignments. The KLP motif, found only in the ancestral-like mouse Ezhip (between residues 530 and 550 in the alignment), is indicated with a green box.

**Supplementary Figure S13.**
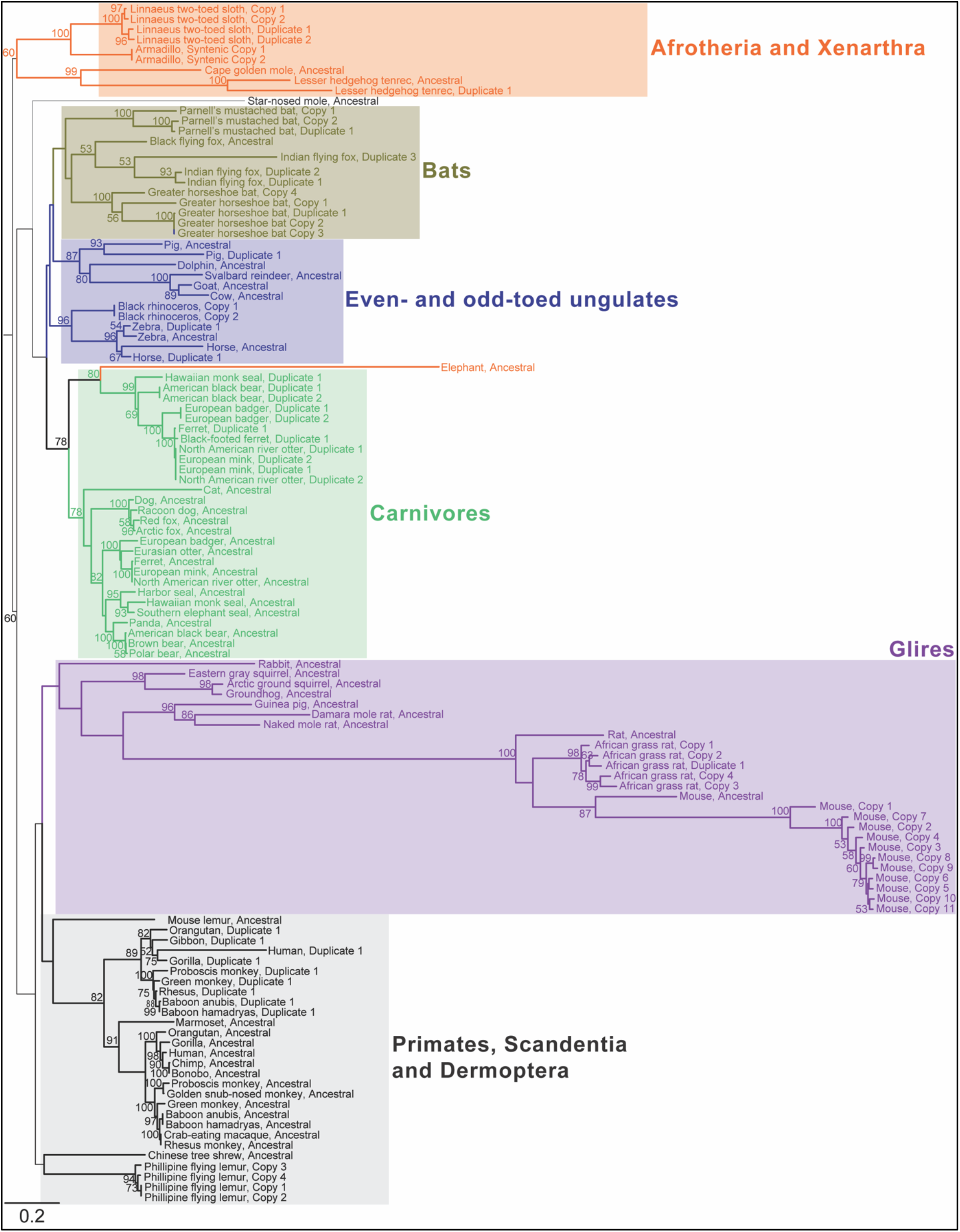
Phylogeny of mammalian EZHIP orthologs and paralogs. Maximum-likelihood protein phylogenetic tree of EZHIP from 70 representative mammalian species (colored as in Figure 1C, Supplementary Data S5). Bootstrap values at selected nodes with >50% support are shown. Names of mammals are indicated at branch tips. In species with multiple copies of EZHIP, ancestral copies identified based on syntenic location are indicated with ‘Ancestral’, syntenic copies that could not be reliably distinguished from ancestral copies are indicated with ‘Copy #’, and duplicate copies that could be reliably distinguished from ancestral EZHIP are indicated with ‘Duplicate #’. The bottom-left scale bar below the phylogeny represents 0.2 substitutions per site.

**Supplementary Figure S14.**
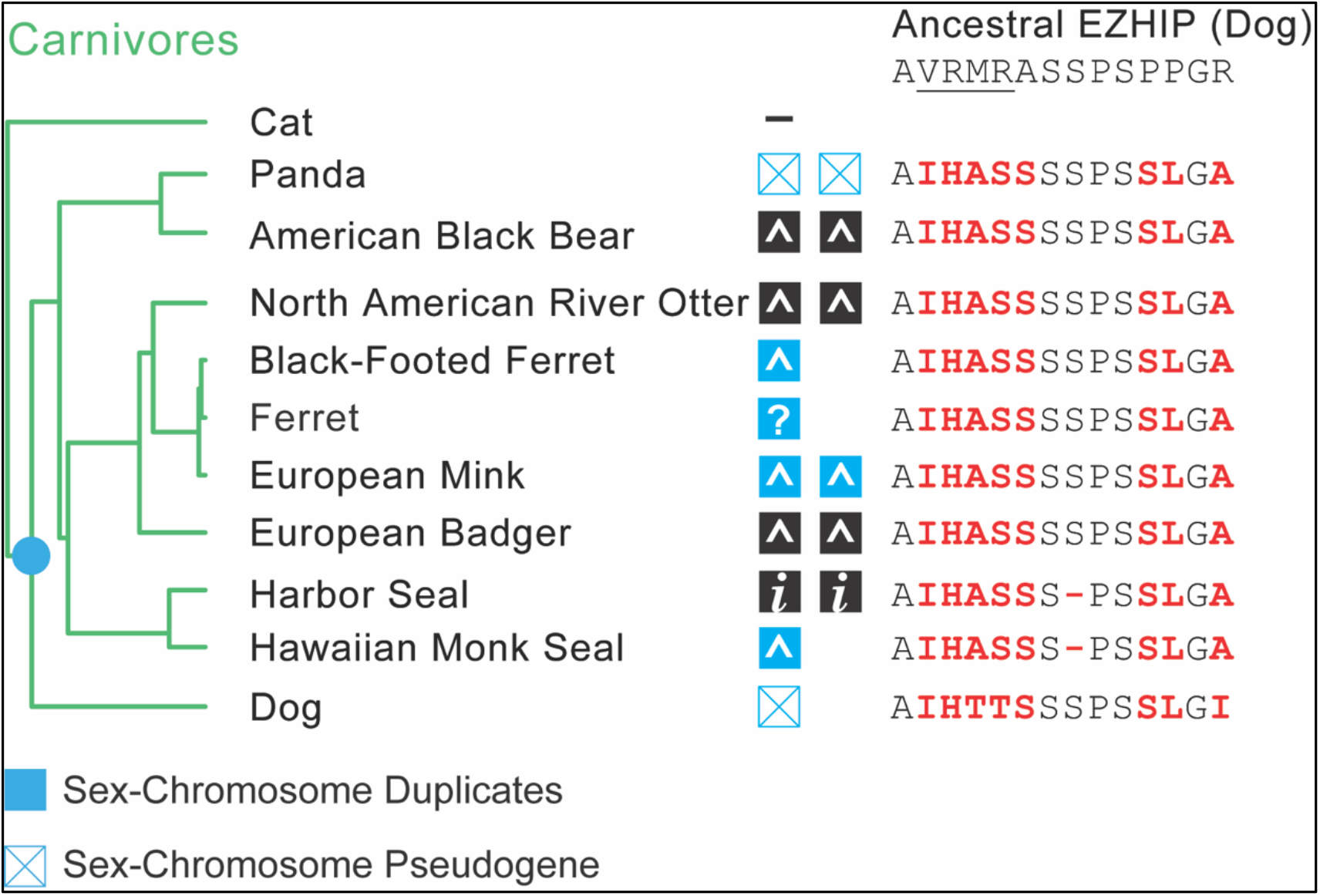
An EZHIP paralog arose in the last common ancestor of carnivores. An EZHIP paralog that can be found in the same syntenic location in carnivores and that has subsequently been duplicated in carnivores is illustrated with boxes alongside a carnivore species tree. Based on synteny across several carnivores and the lack of a duplicate in cat at the syntenic location, we infer that this *EZHIP* paralog arose in the common ancestor of Caniforma (blue dot in species tree). Blue boxes indicate identification on the X chromosome, while black boxes indicate sequences that were not assigned to chromosomes in the genome assembly. Boxes containing an X represent putative pseudogenes, *i* represents incomplete sequences due to gaps in genome assembly, and ^ represents a lack of a KLP sequence in duplicates. Right, the KLP sequence with the oncohistone-mimic region underlined from dog ancestral EZHIP (top) is compared to the same region across the paralogs (one per species), with changes relative to ancestral dog EZHIP in red. Hyphens indicate gaps in the alignment.

**Supplementary Figure S15.**
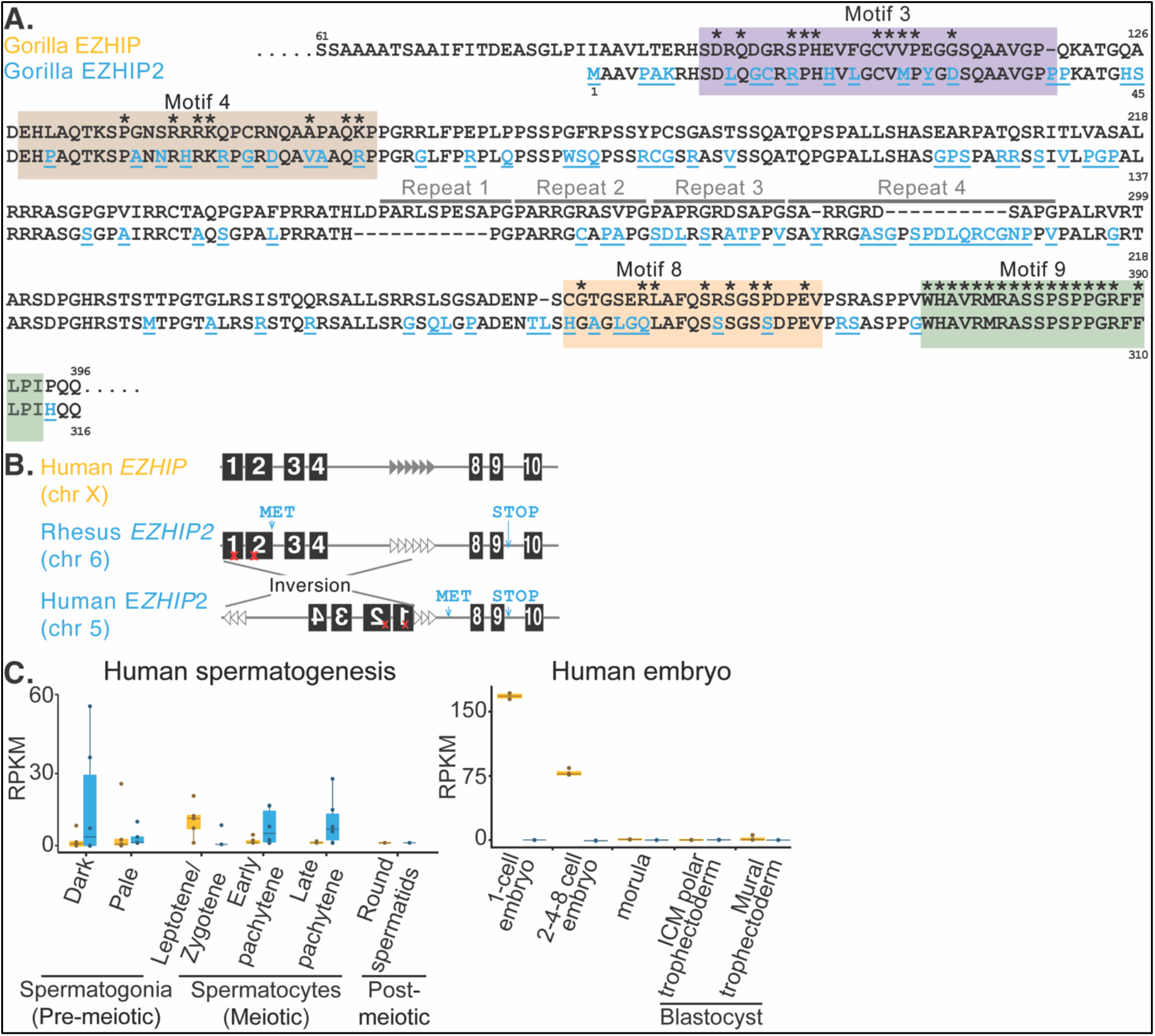
EZHIP and EZHIP2 are similar and are expressed in germ cells. **A.** An alignment of gorilla EZHIP and EZHIP2 is shown with motifs 3, 4, 8, 9 and repeats highlighted. Hyphens indicate gaps in the alignment. Residues that differ between EZHIP and EZHIP2 are colored in blue and underlined. Asterisks indicate residues that we identify as being strongly retained across mammalian EZHIP in Figure 2A. **B.** EZHIP2 may have arisen via retrotransposition and has undergone an inversion in humans. The genomic arrangement of *EZHIP* (503 amino acids) compared to rhesus *EZHIP2*, which encodes a slightly shorter protein (316 amino acids), and human *EZHIP2*, which shows an inversion, but which could still encode an 88 amino acid protein that includes the KLP, is shown. *EZHIP* is on a sex chromosome, but *EZHIP2* is on an autosome at the same syntenic location in the analyzed primates. Red X symbols indicate nonsense mutations in motifs 1 and 2. A methionine was gained after motif 2, and a premature stop codon after motif 9, resulting in a shorter open reading frame for most primate *EZHIP2* sequences. **C**. Expression of *EZHIP* paralogs across stages of human spermatogenesis and embryonic development is plotted as in Figure 4B.

**Supplementary Figure S16.**
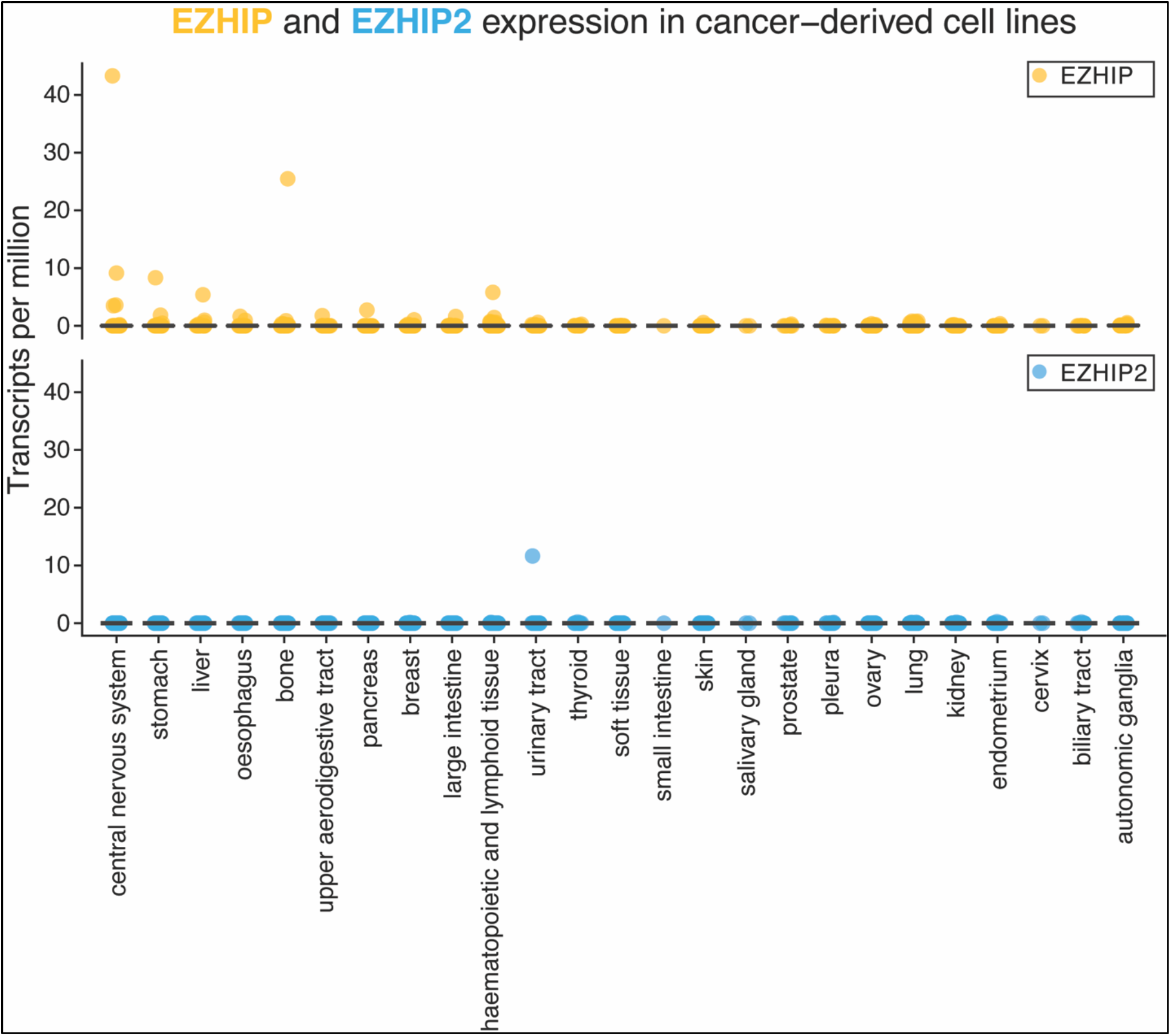
Expression of *EZHIP* and *EZHIP2* across cancer-derived cell lines. Expression of human *EZHIP* (yellow) and *EZHIP2* (blue) mRNA across a variety of cancer-derived cell lines from the Cancer Cell Line Encyclopedia.

**Supplementary Figure S17.**
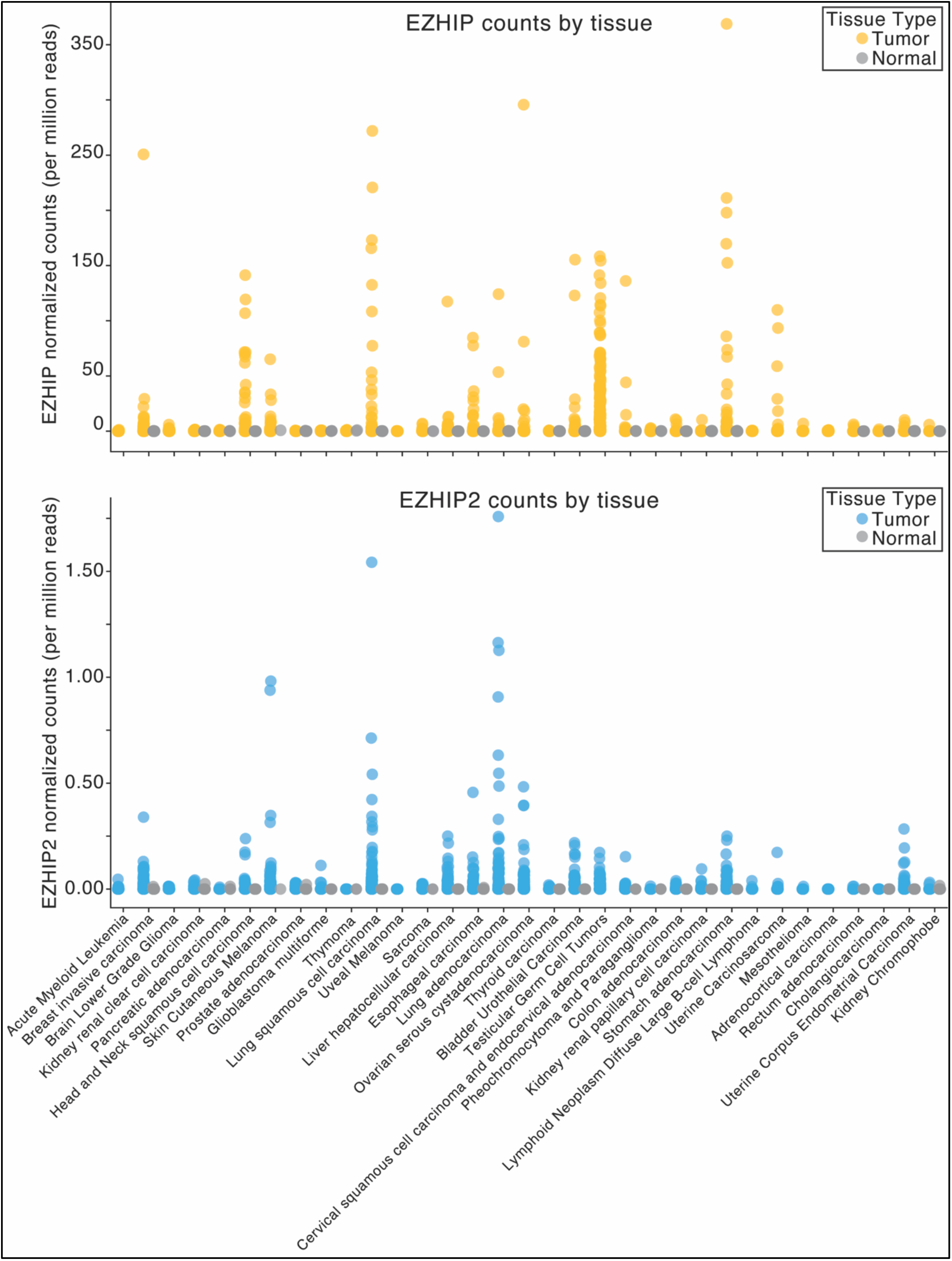
Expression of *EZHIP* and *EZHIP2* across cancer types. Expression of human *EZHIP* and *EZHIP2* is compared between patient-derived normal (gray) and cancer tissues (*EZHIP* in yellow and *EZHIP2* in blue) across a variety of cancer types from The Cancer Genome Atlas (TCGA). Note differences in the Y-axis range of EZHIP and EZHIP2 normalized counts.

**Supplementary Figure S18.**
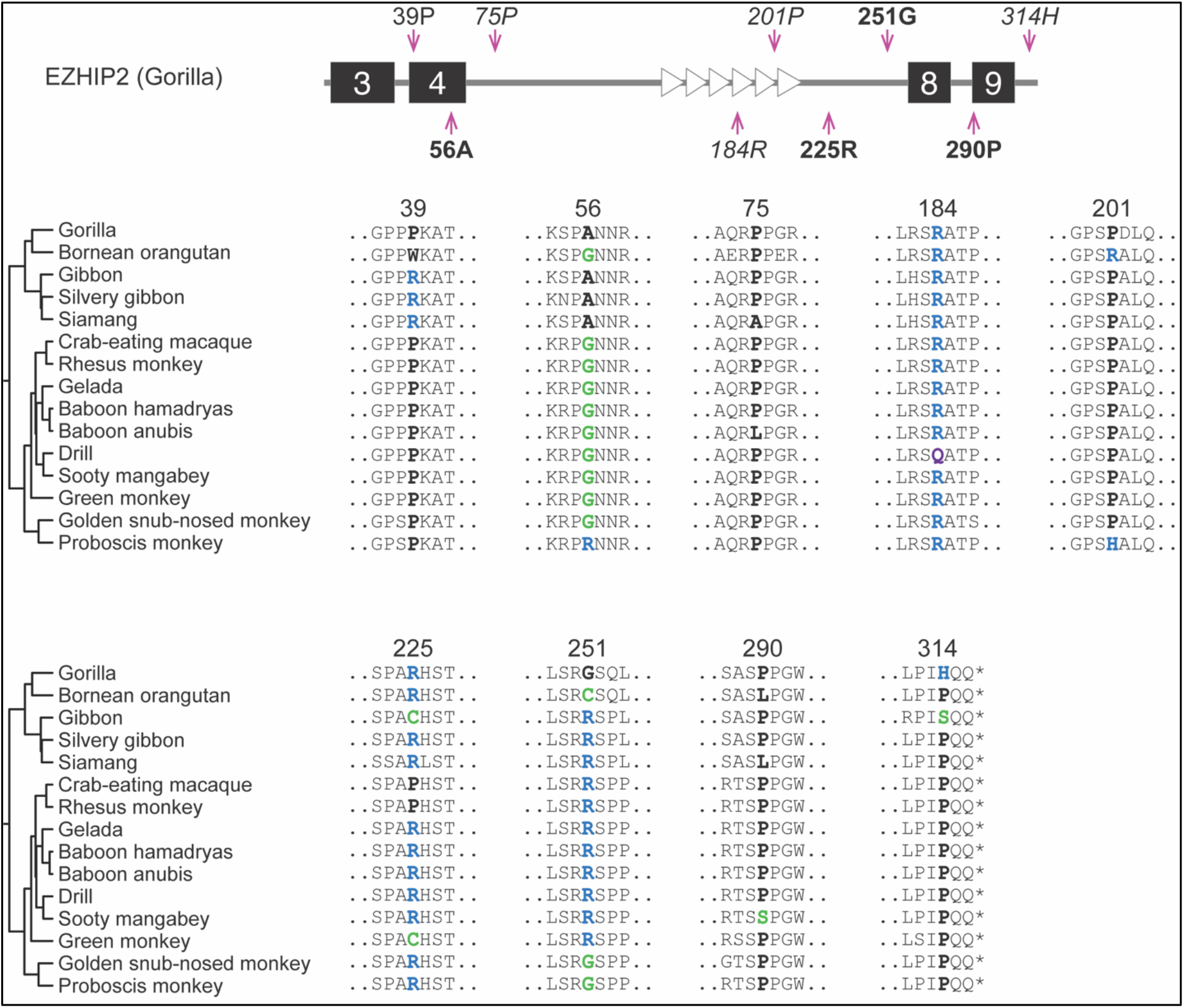
Evidence of positive selection at a subset of sites across simian primate EZHIP2. Alignments of regions surrounding positively selected sites (colored amino residues) in simian primate EZHIP2 identified by PAML and/or FUBAR (also see Fig. 4C). * indicates stop codon. Colors of residues highlight their biochemical properties: hydrophobic (black), negatively charged (blue), positively charged (red), polar (green), and others (purple).

**Table S1:**
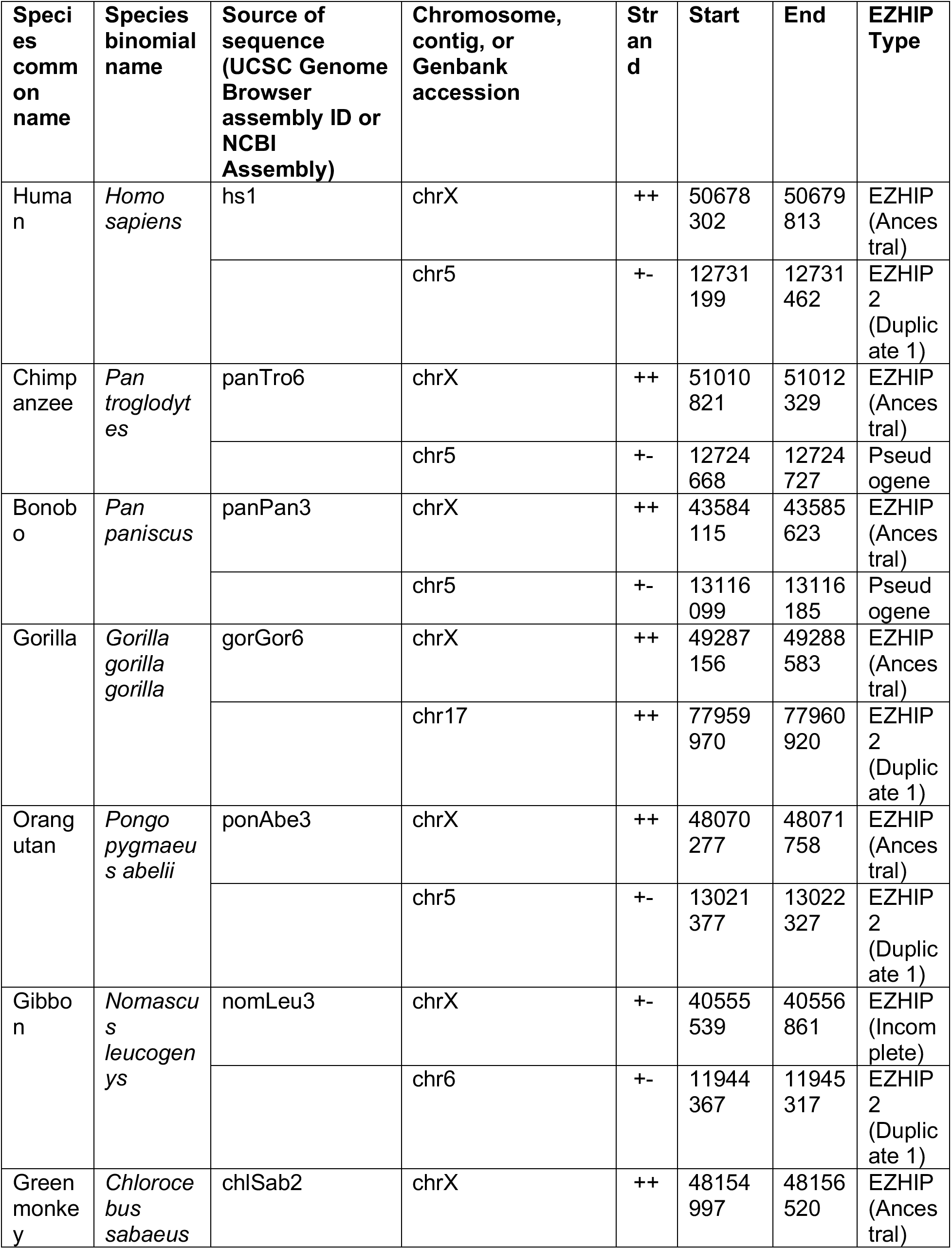
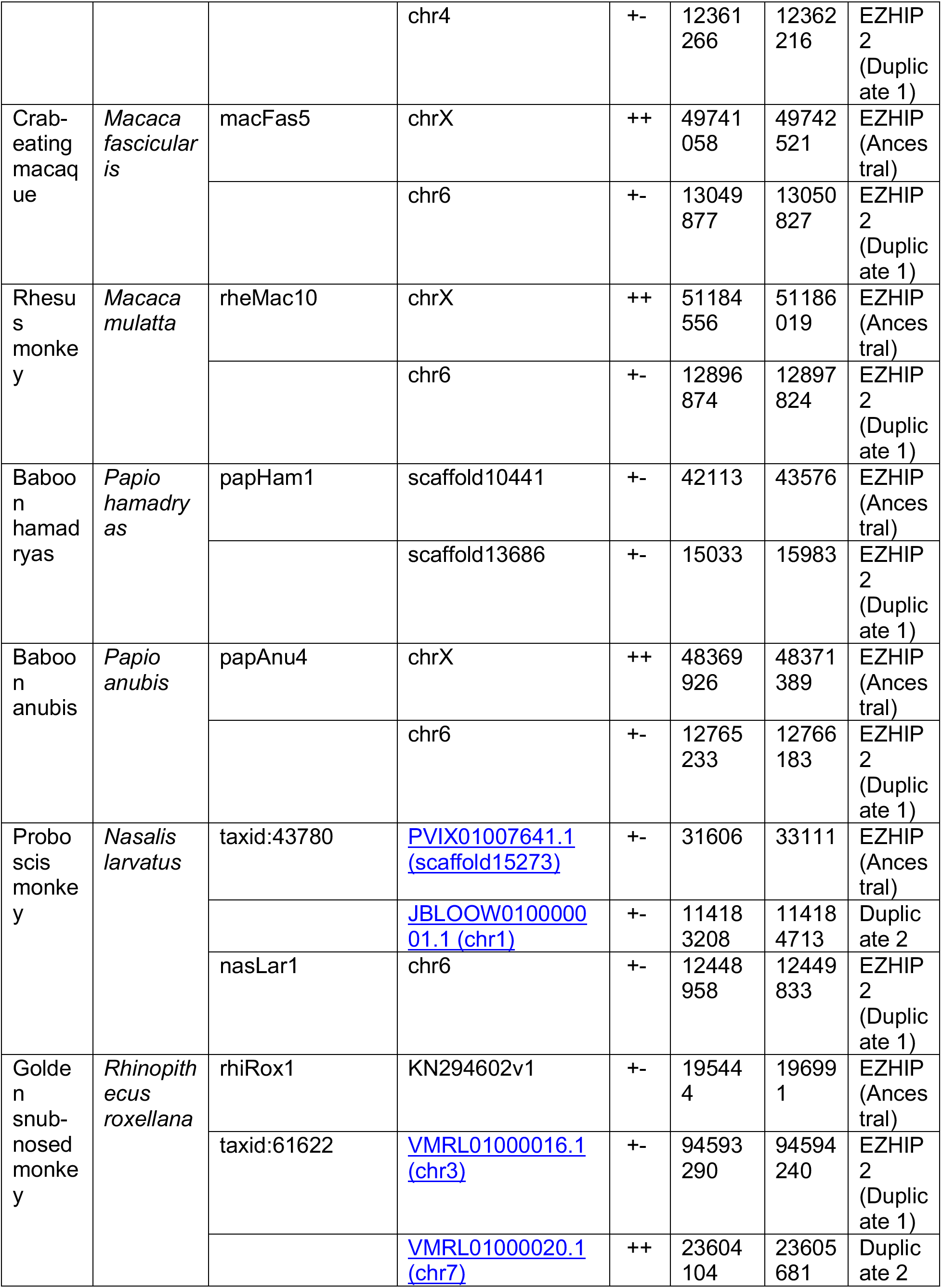
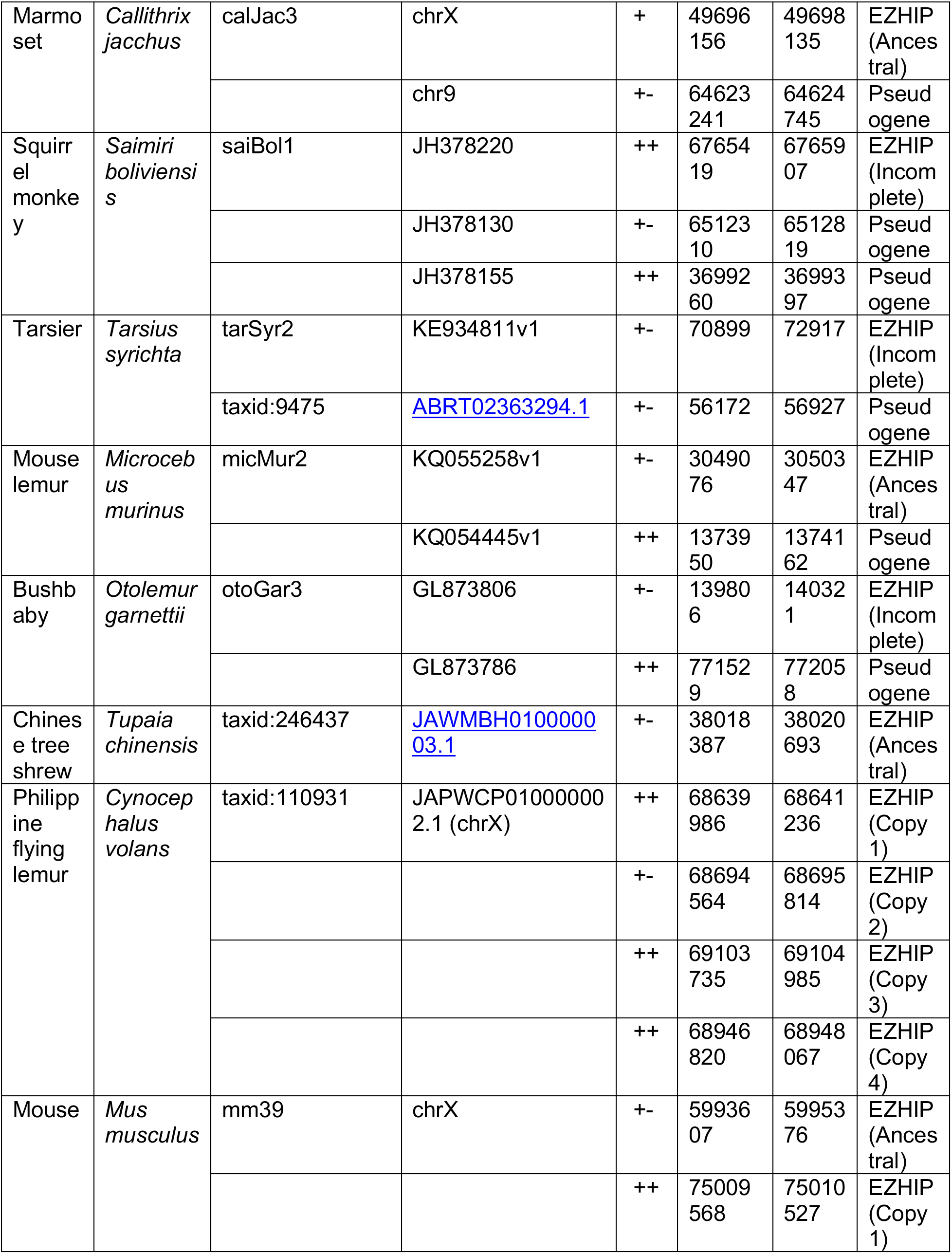
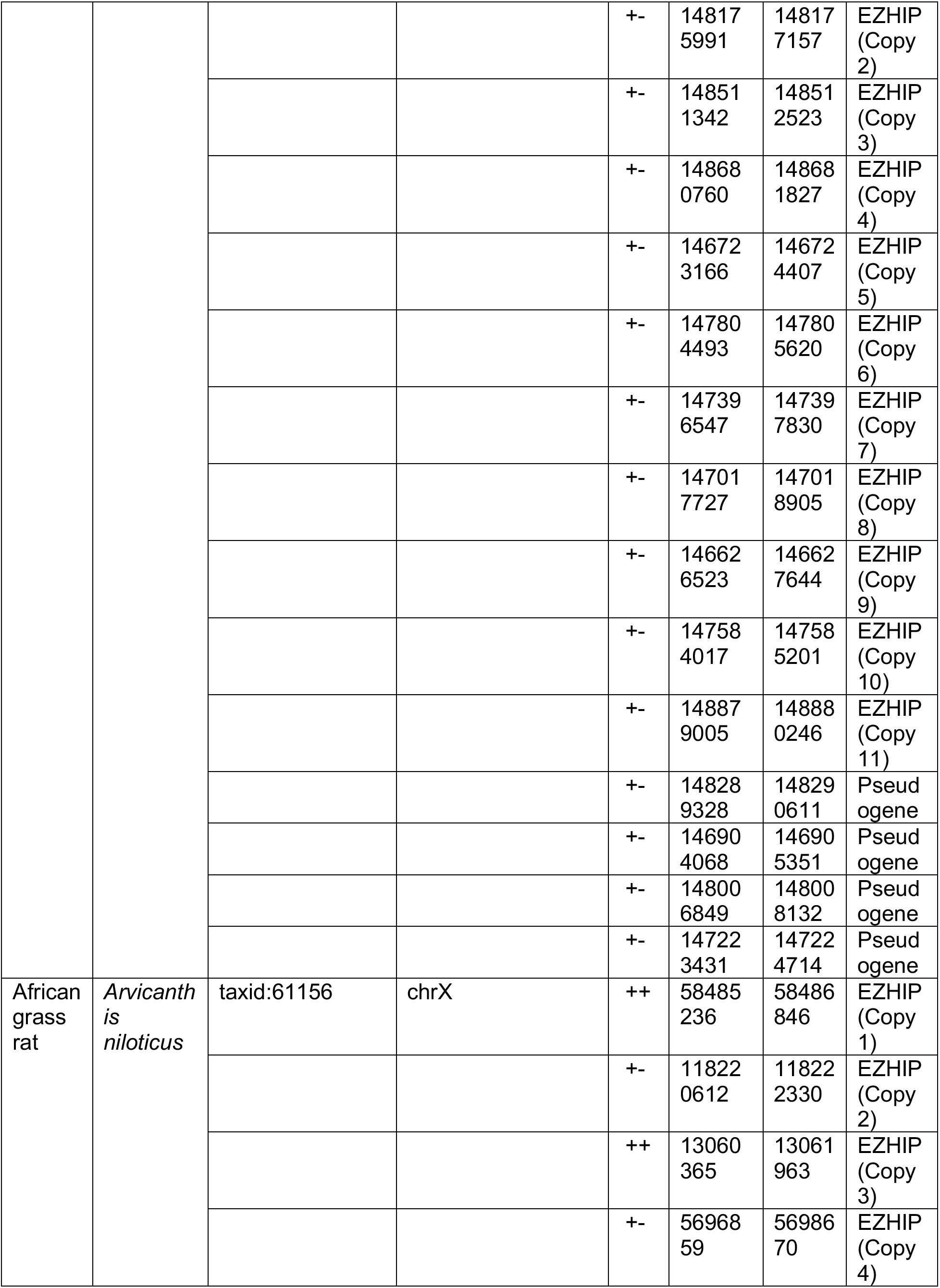
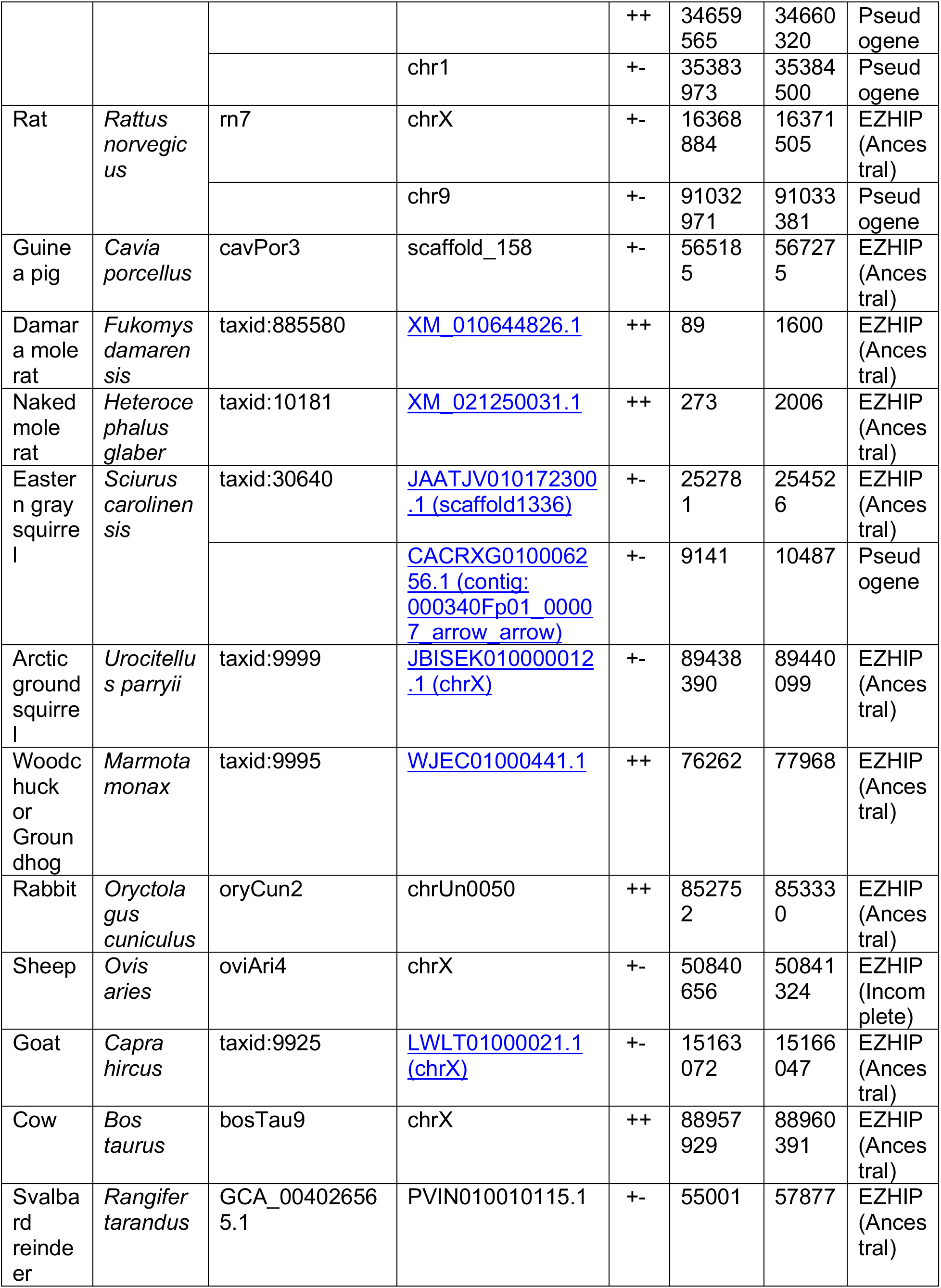
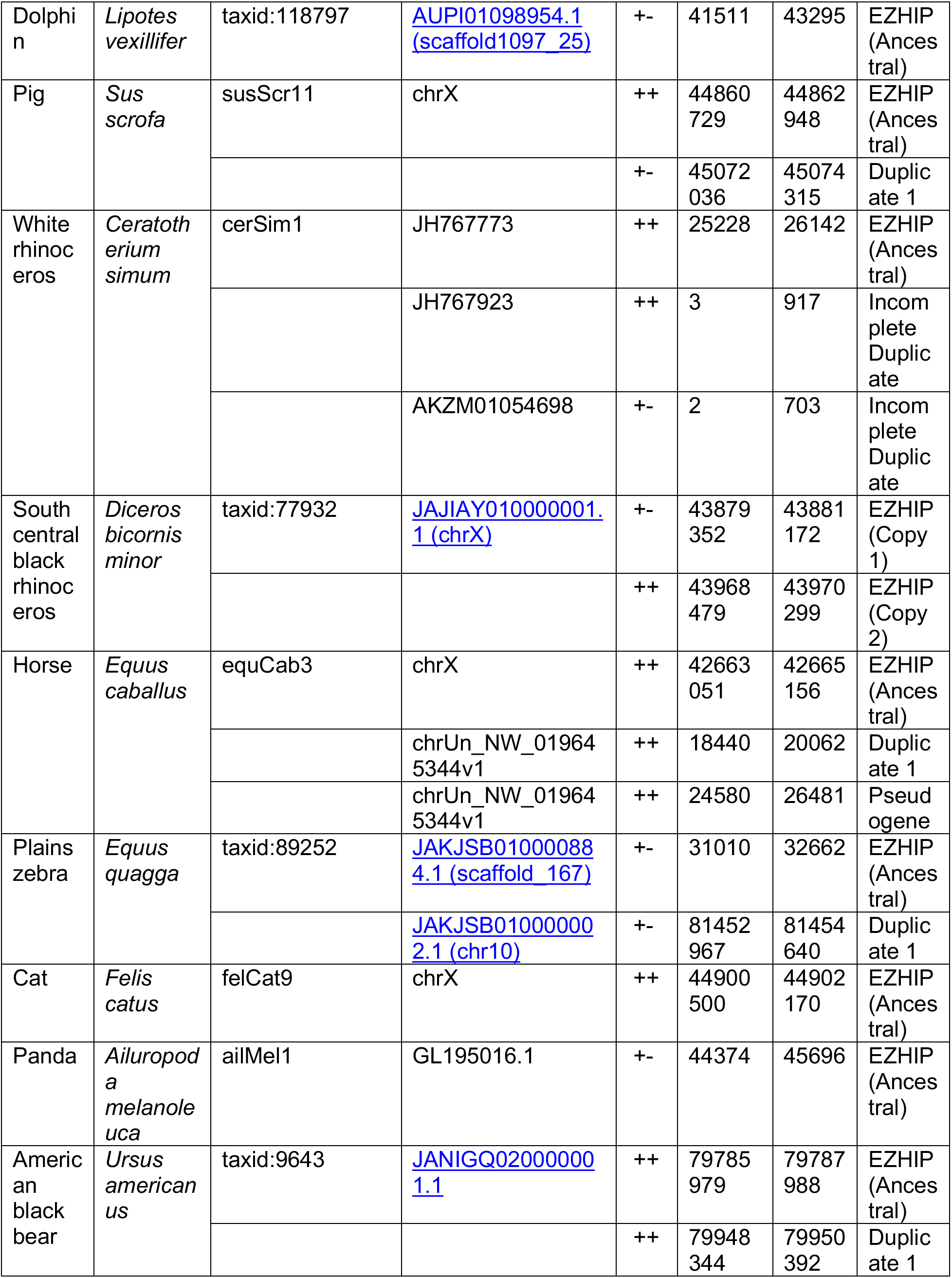
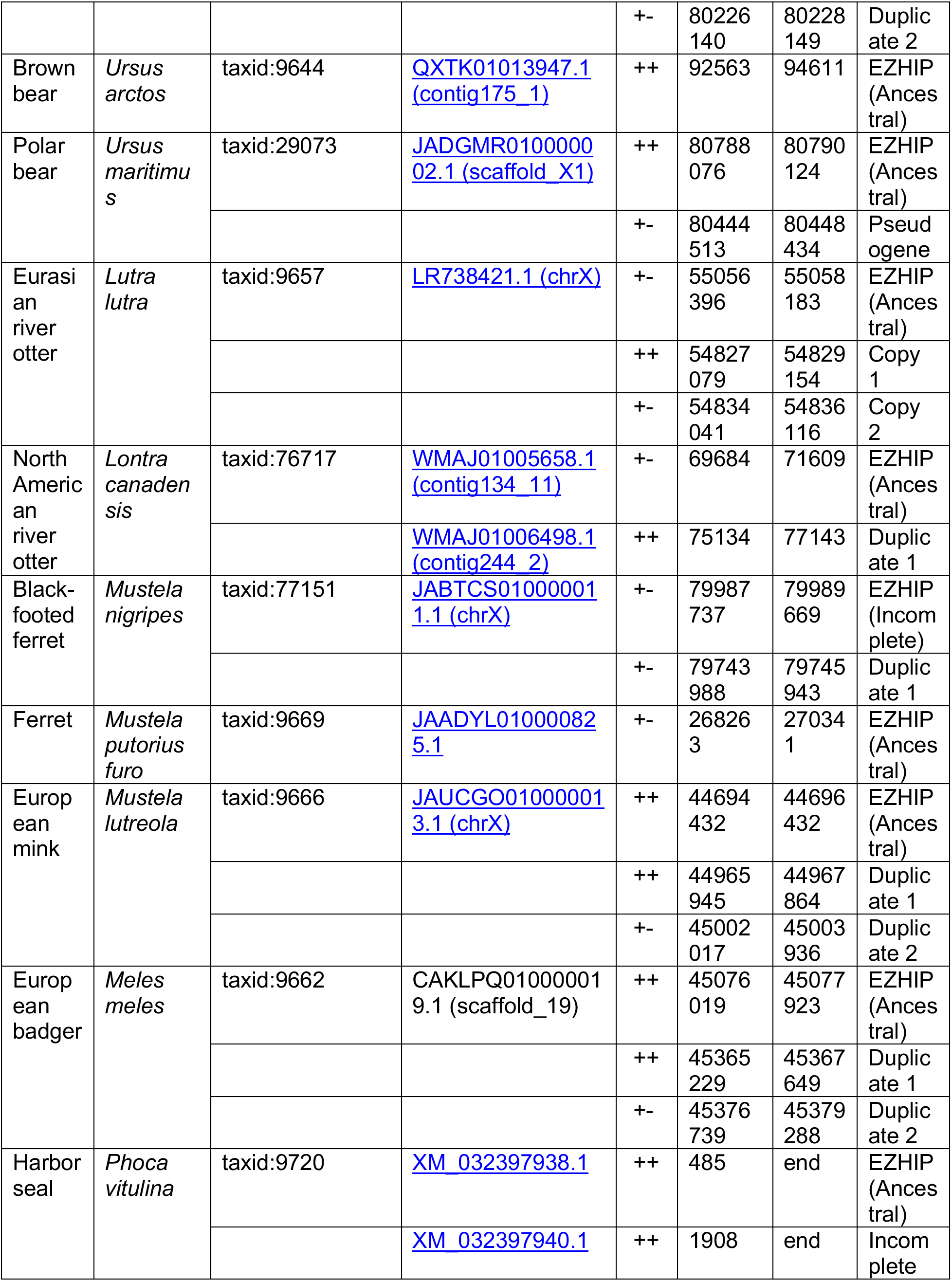
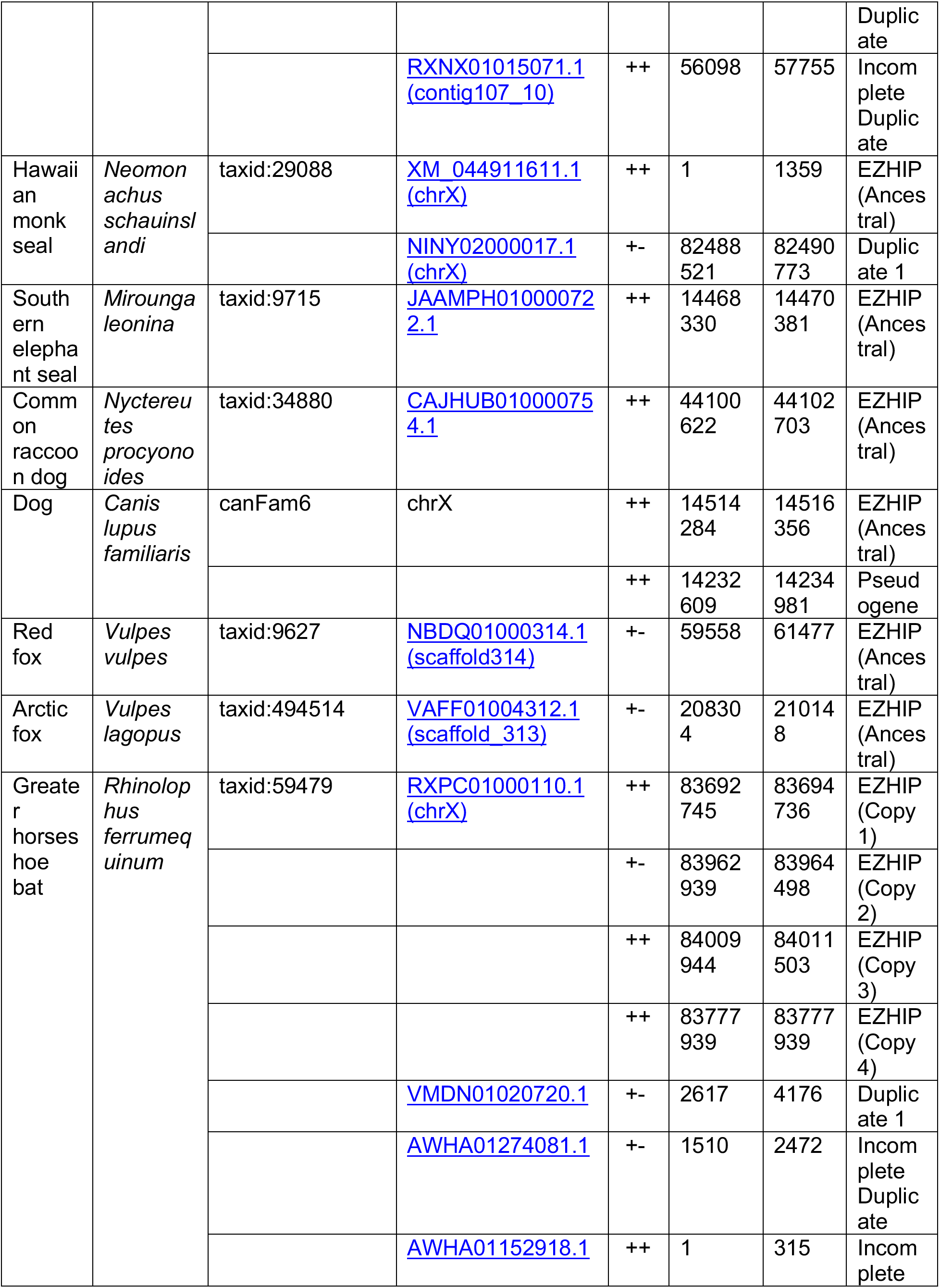
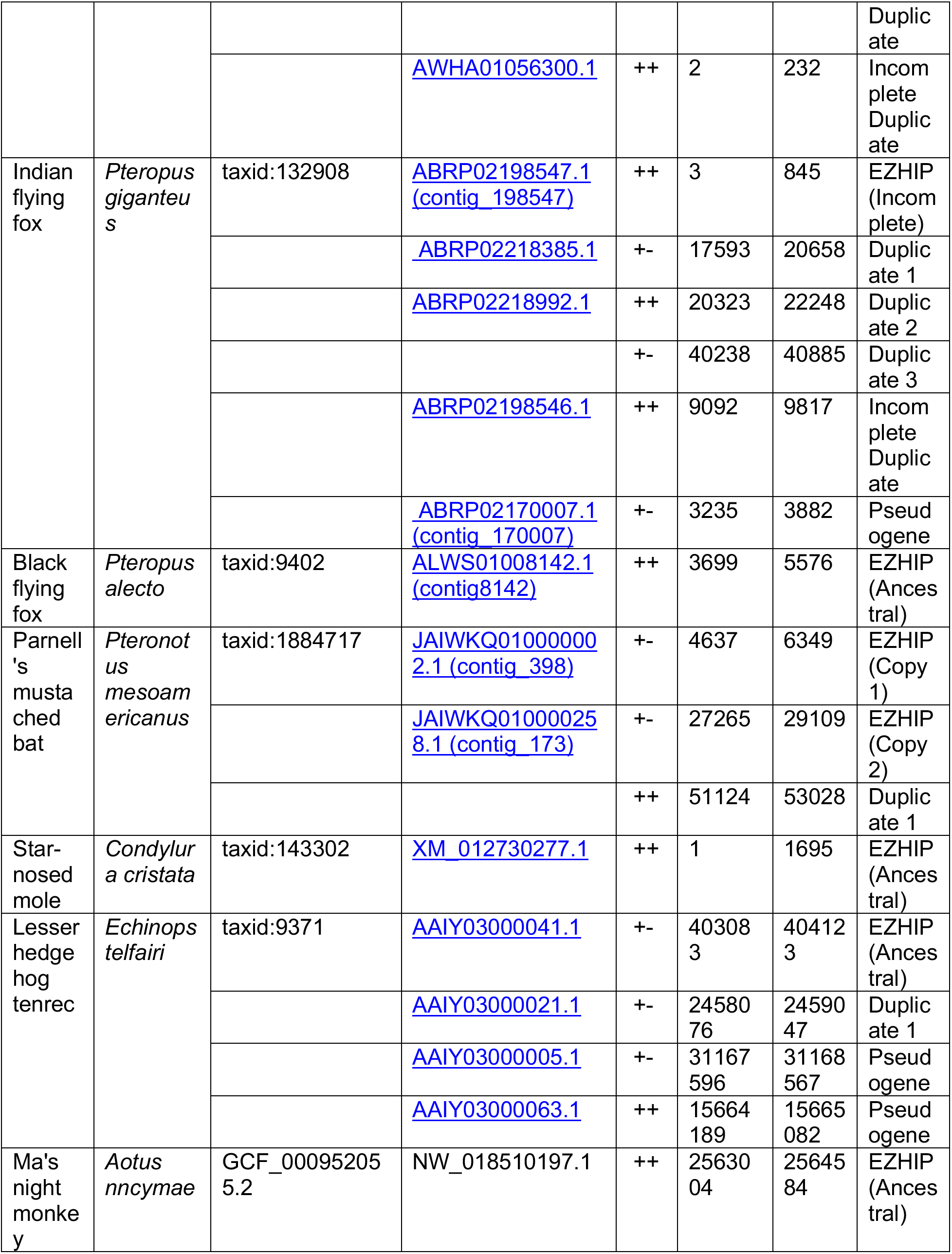
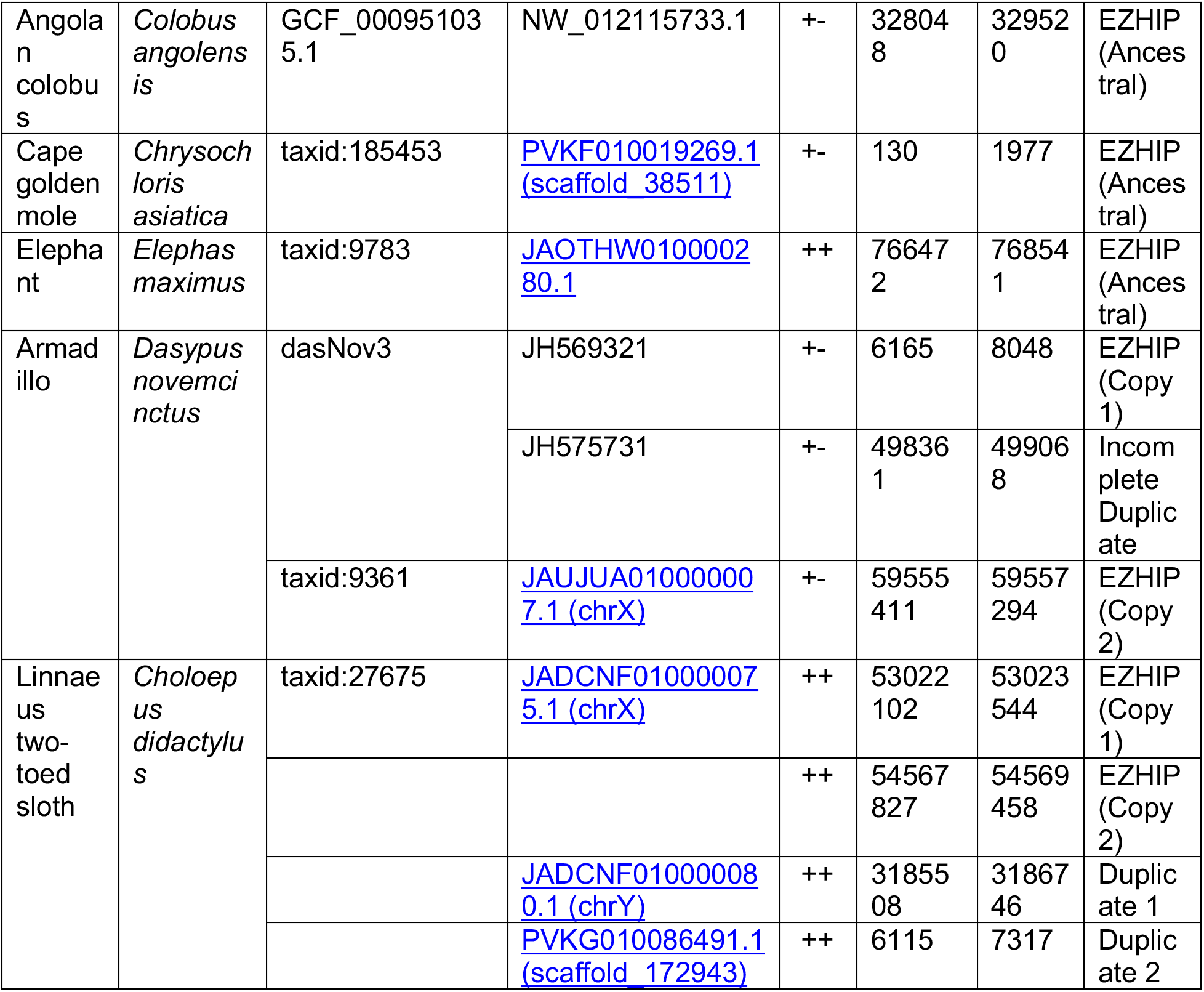
Genomic coordinates of all EZHIP genes in this study.

**Table S2:** Results from purifying selection analyses.

| Gene | Tree file | Num. seqs | Seq. length (nt) | Seq. length (codons) | Codeml selection model | Starting omega | lnL | Num. parameters | 2diffML | Degrees of freedom | p-value | dN/dS |
| --- | --- | --- | --- | --- | --- | --- | --- | --- | --- | --- | --- | --- |
| EZHIP in Afrotheria | Gene Tree | 4 | 999 | 333 | 0 | 0.4 | -3085.830 | 7 |  |  |  | 0.72 |
|  |  |  |  |  | 0, fixed w=1 | 1 | -3090.030 | 6 | 8.3999 | 1 | **0.00375** |  |
| EZHIP in mammals | Gene Tree | 13 | 876 | 292 | 0 | 0.4 | -5964.223 | 25 |  |  |  | 0.85 |
|  |  |  |  |  | 0, fixed w=1 | 1 | -5967.200 | 24 | 5.9531 | 1 | **0.01469** |  |
| EZHIP2 in primates without human | Gene Tree | 17 | 951 | 317 | 0 | 0.4 | -2517.640 | 27 |  |  |  | 1.34 |
|  |  |  |  |  | 0, fixed w=1 | 1 | -2519.428 | 26 | 3.5762 | 1 | **0.05861** |  |

**Table S3:** Divergence of EZHIP in mammalian lineages.

| **EZHIP** |  |  |  |
| --- | --- | --- | --- |
| Species | Divergence time<br>(time tree median<br>time, MYA) | %<br>identity | Time tree range |
| **PRIMATES compared to human** |  |  |  |
| **Human** | 0 | 100 |  |
| Chimp | 6.4 | 96.2 | 6.0 - 6.5 MYA |
| Gorilla | 8.6 | 97.7 | 8.3 - 9.3 MYA |
| Orangutan | 15.2 | 94.7 | 14.5 - 16.3 MYA |
| Baboon (Anubis) | 28.8 | 89.6 | 26.8 - 30.6 MYA |
| Marmoset | 43 | 74.8 | 40.0 - 44.2 MYA |
| Philippine Flying Lemur | 79 | 54.6 | 70.5-86.2 MYA |
| **RODENTS compared to mouse** |  |  |  |
| Mouse | 0 | 100 |  |
| African Grass Rat | 11 | 58.2 | 10.0 - 19.6 MYA |
| Rat | 13.1 | 34.5 | 12.2 - 14.2 MYA |
| Damara mole-rat | 70 | 37.7 | 56.1 - 77.0 MYA |
| Groundhog | 69 | 34.4 | 64.4 - 74.0 MYA |
| **TWO-TOED UNGULATES compared to goat** |  |  |  |
| **Goat** | 0 | 100 |  |
| Cow | 24.6 | 86.2 | 19.1 - 30.8 MYA |
| Reindeer | 22.8 | 80.1 | 18.5 - 27.8 MYA |
| Pig | 62 | 59.6 | 52.0 - 63.9 MYA |
| Black Rhino | 76 | 59.8 | 72.8 - 78.3 MYA |
| Horse | 76 | 53.4 | 72.8 - 78.3 MYA |
| Zebra | 76 | 57 | 72.8 - 78.3 MYA |
| **CARNIVORES compared to dog** |  |  |  |
| **Dog** | 0 | 100 |  |
| Red fox | 12.2 | 94 | 6.6 - 17.1 MYA |
| Brown bear | 45 | 77.6 | 43.4 - 48.5 MYA |
| Northe American River otter | 45 | 74.9 | 43.4 - 48.5 MYA |
| Cat | 55 | 63.9 | 52.9 - 57.3 MYA |
| Panda | 1530 | 79.4 | 1085.0 - 1671.0 MYA |

| **EZH2** |  |  |  |
| --- | --- | --- | --- |
| Species | Divergence time (time<br>tree median time, MYA) | % identity | Time tree range |
| **PRIMATES compared to human** |  |  |  |
| **Human** | 0 | 100 |  |
| Chimp | 6.4 | 92.5 | 6.0 - 6.5 MYA |
| Gorilla | 8.6 | 99.1 | 8.3 - 9.3 MYA |
| Orangutan | 15.2 | 100 | 14.5 - 16.3 MYA |
| Baboon (Anubis) | 28.8 | 100 | 26.8 - 30.6 MYA |
| Marmoset | 43 | 92.9 | 40.0 - 44.2 MYA |
| Phillipine Flying Lemur | 79 | 98.8 | 70.5-86.2 MYA |
| **RODENTS compared to mouse** |  |  |  |
| Mouse | 0 | 100 |  |
| African Grass Rat | 11 | 99.5 | 10.0 - 19.6 MYA |
| Rat | 13.1 | 99.7 | 12.2 - 14.2 MYA |
| Damara mole-rat | 70 | 98.5 | 56.1 - 77.0 MYA |
| Groundhog | 69 | 98.5 | 64.4 - 74.0 MYA |

**Table S4:** Detailed table of results from all analyses of positive selection (simian primates)

| Alignme<br>nt | Species<br>group | T<br>r<br>e<br>e<br>f<br>i<br>l<br>e | Num<br>.<br>seqs | Seq.<br>lengt<br>h (nt) | Seq.<br>length<br>(codon<br>s) | Codeml<br>selecti<br>on<br>model | Startin<br>g<br>omega | Codo<br>n<br>mode<br>l | InL | Nu<br>m.<br>par<br>am<br>ete<br>rs | Test | 2diffML | Degr<br>ees<br>of<br>freed<br>om | p-value | % sites<br>under<br>positiv<br>e<br>selecti<br>on | dN/dS<br>of sites<br>under<br>positiv<br>e<br>selecti<br>on | FUBAR<br>positively<br>selected<br>sites<br>(Posterior<br>probability>0<br>.9). Bold<br>residues are<br>common with<br>PAML<br>analyses |
| --- | --- | --- | --- | --- | --- | --- | --- | --- | --- | --- | --- | --- | --- | --- | --- | --- | --- |
| EZHIP<br>without<br>repeats | Simian<br>Primates | G<br>e<br>n<br>e<br>T<br>r<br>e<br>e | 25 | 1245 | 415 | 7 | 0.4 | 2 | -4442 | 36 |  |  |  |  |  |  | 93L, 158P,<br>171P, 186C,<br>348R, 377E in<br>human EZHIP |
|  |  |  |  |  |  | 8 |  |  | -<br>4420.6 | 38 | M7vsM8 | 42.79 | 2 | 5.1E-10 | 12.2 | 4.08 |  |
|  |  |  |  |  |  | 8a |  |  | -<br>4435.7 | 37 | M8vsM8a | 30.21 | 1 | 3.9E-08 |  |  |  |
| EZHIP2<br>without<br>human | Simian<br>Primates | G<br>e<br>n<br>e<br>T<br>r<br>e<br>e | 17 | 951 | 317 | 7 | 0.4 | 2 | -<br>2519.4 | 28 |  |  |  |  |  |  | 56A, 75P,<br>184R, 201P<br>225R, 251G,<br>290P, 314H in<br>Gorilla EZHIP |
|  |  |  |  |  |  | 8 |  |  | -<br>2513.3 | 30 | M7vsM8 | 12.28 | 2 | 0.00215 | 11.30 | 4.97 |  |
|  |  |  |  |  |  | 8a |  |  | -<br>2519.4 | 29 | M8vsM8a | 12.28 | 1 | 0.00046 |  |  |  |
| EZH2 | Simian<br>Primates | G<br>e<br>n<br>e<br>T<br>r<br>e<br>e | 25 | 1245 | 415 | 7 | 0.4 | 2 | -4442 | 36 |  |  |  |  |  |  | 93L, 158P,<br>171P, 186C,<br>348R, 377E in<br>human EZHIP |
|  |  |  |  |  |  | 8 |  |  | -<br>4420.6 | 38 | M7vsM8 | 42.79 | 2 | 5.1E-10 | 12.2 | 4.08 |  |
|  |  |  |  |  |  | 8a |  |  | -<br>4435.7 | 37 | M8vsM8a | 30.21 | 1 | 3.9E-08 |  |  |  |

**Table S5:** List of SRA datasets used for RNA-seq analysis.

| Figure | Species | SRA run ID | Sample type | Data source |
| --- | --- | --- | --- | --- |
| Figure 4B | Human | SRR306838 | brain | Brawand et al, 2011 |
|  |  | SRR306839 | brain | Brawand et al, 2011 |
|  |  | SRR306840 | brain | Brawand et al, 2011 |
|  |  | SRR306841 | brain | Brawand et al, 2011 |
|  |  | SRR306842 | brain | Brawand et al, 2011 |
|  |  | SRR306843 | brain | Brawand et al, 2011 |
|  |  | SRR306847 | heart | Brawand et al, 2011 |
|  |  | SRR306848 | heart | Brawand et al, 2011 |
|  |  | SRR306849 | heart | Brawand et al, 2011 |
|  |  | SRR306850 | heart | Brawand et al, 2011 |
|  |  | SRR306851 | kidney | Brawand et al, 2011 |
|  |  | SRR306852 | kidney | Brawand et al, 2011 |
|  |  | SRR306853 | kidney | Brawand et al, 2011 |
|  |  | SRR306854 | liver | Brawand et al, 2011 |
|  |  | SRR306855 | liver | Brawand et al, 2011 |
|  |  | SRR306856 | liver | Brawand et al, 2011 |
|  |  | SRR306857 | testis | Brawand et al, 2011 |
|  |  | SRR306858 | testis | Brawand et al, 2011 |
|  |  | SRR2156552 | ovary | Williams et al, 2015 |
|  |  | SRR2156551 | ovary | Williams et al, 2015 |
|  |  | SRR2156550 | ovary | Williams et al, 2015 |
| Figure 4B | Rhesus macaque | SRR306777 | brain | Brawand et al, 2011 |
|  |  | SRR306778 | brain | Brawand et al, 2011 |
|  |  | SRR306779 | brain | Brawand et al, 2011 |
|  |  | SRR306782 | heart | Brawand et al, 2011 |
|  |  | SRR306783 | heart | Brawand et al, 2011 |
|  |  | SRR306784 | kidney | Brawand et al, 2011 |
|  |  | SRR306785 | kidney | Brawand et al, 2011 |
|  |  | SRR306786 | liver | Brawand et al, 2011 |
|  |  | SRR306787 | liver | Brawand et al, 2011 |
|  |  | SRR306788 | liver | Brawand et al, 2011 |
|  |  | SRR306789 | testis | Brawand et al, 2011 |
|  |  | SRR306790 | testis | Brawand et al, 2011 |
| Figure S13C | Human | SRR3146452 | Dark spermatogonia | Jan et al. 2017 |
|  |  | SRR6726012 | Dark spermatogonia | Jan et al. 2017 |
|  |  | SRR6726013 | Dark spermatogonia | Jan et al. 2017 |
|  |  | SRR6726014 | Dark spermatogonia | Jan et al. 2017 |
|  |  | SRR6726015 | Dark spermatogonia | Jan et al. 2017 |
|  |  | SRR6726016 | Dark spermatogonia | Jan et al. 2017 |
|  |  | SRR3146494 | Pale spermatogonia | Jan et al. 2017 |
|  |  | SRR6726159 | Pale spermatogonia | Jan et al. 2017 |
|  |  | SRR6726160 | Pale spermatogonia | Jan et al. 2017 |
|  |  | SRR6726161 | Pale spermatogonia | Jan et al. 2017 |
|  |  | SRR6726162 | Pale spermatogonia | Jan et al. 2017 |
|  |  | SRR3146504 | Leptotene/zygotene spermatocytes | Jan et al. 2017 |
|  |  | SRR6726087 | Leptotene/zygotene spermatocytes | Jan et al. 2017 |
|  |  | SRR6726088 | Leptotene/zygotene spermatocytes | Jan et al. 2017 |
|  |  | SRR6726089 | Leptotene/zygotene spermatocytes | Jan et al. 2017 |
|  |  | SRR6726090 | Leptotene/zygotene spermatocytes | Jan et al. 2017 |
|  |  | SRR3146497 | Early pachytene spermatocytes | Jan et al. 2017 |
|  |  | SRR6726153 | Early pachytene spermatocytes | Jan et al. 2017 |
|  |  | SRR6726154 | Early pachytene spermatocytes | Jan et al. 2017 |
|  |  | SRR6726155 | Early pachytene spermatocytes | Jan et al. 2017 |
|  |  | SRR6726156 | Early pachytene spermatocytes | Jan et al. 2017 |
|  |  | SRR6726157 | Early pachytene spermatocytes | Jan et al. 2017 |
|  |  | SRR3146507 | Late pachytene spermatocytes | Jan et al. 2017 |
|  |  | SRR6726042 | Late pachytene spermatocytes | Jan et al. 2017 |
|  |  | SRR6726043 | Late pachytene spermatocytes | Jan et al. 2017 |
|  |  | SRR6726044 | Late pachytene spermatocytes | Jan et al. 2017 |
|  |  | SRR6726045 | Late pachytene spermatocytes | Jan et al. 2017 |
|  |  | SRR6726046 | Late pachytene spermatocytes | Jan et al. 2017 |
|  |  | SRR3146508 | Round spermatids | Jan et al. 2017 |
|  |  | SRR6726017 | Round spermatids | Jan et al. 2017 |
|  |  | SRR6726018 | Round spermatids | Jan et al. 2017 |
|  |  | SRR6726019 | Round spermatids | Jan et al. 2017 |
|  |  | SRR6726020 | Round spermatids | Jan et al. 2017 |
|  |  | SRR6726021 | Round spermatids | Jan et al. 2017 |
| Figure S13C | Human | SRR2183367 | 1-cell embryo | Hendrickson et al. 2017 |
|  |  | SRR2183368 | 1-cell embryo | Hendrickson et al. 2017 |
|  |  | SRR2183369 | 1-cell embryo | Hendrickson et al. 2017 |
|  |  | SRR2183370 | 1-cell embryo | Hendrickson et al. 2017 |
|  |  | SRR2183371 | 2-4-8 cell embryo | Hendrickson et al. 2017 |
|  |  | SRR2183372 | 2-4-8 cell embryo | Hendrickson et al. 2017 |
|  |  | SRR2183373 | 2-4-8 cell embryo | Hendrickson et al. 2017 |
|  |  | SRR2183374 | 2-4-8 cell embryo | Hendrickson et al. 2017 |
|  |  | SRR2183375 | Morula | Hendrickson et al. 2017 |
|  |  | SRR2183376 | Morula | Hendrickson et al. 2017 |
|  |  | SRR2183377 | Morula | Hendrickson et al. 2017 |
|  |  | SRR2183378 | Morula | Hendrickson et al. 2017 |
|  |  | SRR2183379 | ICM polar trophectoderm | Hendrickson et al. 2017 |
|  |  | SRR2183380 | ICM polar trophectoderm | Hendrickson et al. 2017 |
|  |  | SRR2183381 | ICM polar trophectoderm | Hendrickson et al. 2017 |
|  |  | SRR2183382 | ICM polar trophectoderm | Hendrickson et al. 2017 |
|  |  | SRR2183383 | mural trophectoderm | Hendrickson et al. 2017 |
|  |  | SRR2183384 | mural trophectoderm | Hendrickson et al. 2017 |
|  |  | SRR2183385 | mural trophectoderm | Hendrickson et al. 2017 |
|  |  | SRR2183386 | mural trophectoderm | Hendrickson et al. 2017 |

